# What is the best method for estimating ancestral states from discrete characters?

**DOI:** 10.1101/2023.08.31.555762

**Authors:** Joseph N Keating

## Abstract

Ancestral state estimation is a formal phylogenetic method for inferring the nature of ancestors and performing tests of character evolution. As such, it is among the most important tools available to evolutionary biologists. However, there are a profusion of methods available, the accuracy of which remains unclear. Here I use a simulation approach to test between parsimony and likelihood methods for estimating ancestral states from discrete binary characters. I simulate 500 characters using 15 different Markov generating models, a range of tree sizes (8-256 tips) and three topologies representing end members of tree symmetry and branch length heterogeneity. Simulated tip states were subjected to ancestral state estimation under the Equal Rates (ER) and All-Rates-Different (ARD) models, as well as under parsimony assuming accelerated transformations (ACCTRAN). The results demonstrate that both parsimony and likelihood approaches obtain high accuracy applied to trees with more tips. Parsimony performs poorly when trees contain long branches, whereas the ER model performs well across simulations and is reasonably robust to model violation. The ER model frequently outperforms the ARD model, even when data are simulated using unequal rates. Furthermore, the ER model exhibits less transition rate error when compared to ER models. These results suggest that ARD models may be overparameterized when character data is limited. Surprisingly, the difference in likelihood-based information criteria between models was found to be a poor predictor of difference in model error; better fitting models are not necessarily more accurate. However, there is a strong correlation between model uncertainty and model error; likelihood models with more certain ancestral state estimates are typically more accurate. Using empirical morphological datasets, I demonstrate that applying different methods often results in substantively different ancestral state estimates. The results of the simulation study highlight the importance of incorporating fossils in ancestral state estimation. Fossils increase the total number of tips, break long branches and are closer to internal nodes, thereby lowering average branch length and overall branch length heterogeneity of trees. These factors will all contribute to increasing the accuracy of ancestral state estimates, irrespective of the method used.

## Introduction

Ancestral state estimation encompasses a diverse set of formalised methods aimed at inferring the evolutionary history of a character from observed character states in a sample of species, as well as knowledge of the evolutionary relationships of those species. Character data can be continuous such as body size (e.g. Pimiento, C., Cantalapiedra, J.L., et al. 2019, Moon, B.C. and Stubbs, T.L. 2020), basal metabolic rate (e.g. Legendre, L.J., Guénard, G., et al. 2016) or bite force (e.g. Sakamoto, M., Ruta, M., et al. 2019). Alternatively, characters can be discrete (i.e. absence/presence or condition of a character). Discrete ancestral state estimation is an exceptionally versatile tool that plays a pivotal role in understanding the evolution of various biological phenomena including habitat/niche preferences (e.g. Sallan, L., Friedman, M., et al. 2018), hosts-parasite/pathogen interactions (e.g. Ploch, S., Kruse, J., et al. 2022), behaviour (e.g. Ocampo, D., De Silva, T.N., et al. 2023), morphology (e.g. Sokoloff, D.D., Remizowa, M.V., et al. 2018, Gai, Z., Li, Q., et al. 2022) and developmental processes (e.g. Bonett, R.M., Steffen, M.A., et al. 2014, Jiang, B., He, Y., et al. 2023). Moreover, its potential extends beyond biological studies, reaching into domains such as linguistics, social sciences, and humanities, where it can provide insights into the evolution of language (e.g. Sheard, C., Bowern, C., et al. 2020), marriage types (e.g. Walker, R.S., Hill, K.R., et al. 2011), religion (e.g. Peoples, H.C., Duda, P., et al. 2016, Watts, J., Sheehan, O., et al. 2016), and more.

Discrete characters are typically analysed using parsimony or likelihood approaches (e.g. Maximum likelihood, Bayesian Inference). Parsimony methods optimise the solution which requires the fewest character state changes for the specified tree topology. These methods do not typically utilise information from branch lengths. Likelihood approaches, on the other hand, optimise solutions based on a specified tree; set of branch lengths; and model of character evolution (Joy, J.B., Liang, R.H., et al. 2016, Harmon, L. 2018). Discrete character evolution is typically modelled using a continuous time Markov process, which is specified using a transition rate matrix describing the instantaneous probabilities of transitions between states. The simplest model of discrete character evolution is the Equal Rates (ER) model, which describes transitions between any state using a single rate parameter (i.e. all state transitions occur with equal probability). Relaxing the assumptions of the ER model allows for more complex models of evolution. For example, the All-Rates-Different (ARD) model permits each state transition to occur with a unique probability (Harmon, L. 2018).

Although conceptually simple, accurately estimating the ancestral states of discrete characters poses a significant challenge, primarily because a single character and tree topology contain little information from which to infer evolutionary history. (Schluter, D., Price, T., et al. 1997, Cunningham, C.W., Omland, K.E., et al. 1998, Cunningham, C.W. 1999, Mooers, A.Ø. and Schluter, D. 1999, Gascuel, O. and Steel, M. 2020, Holland, B.R., Ketelaar-Jones, S., et al. 2020, Boyko, J.D. and Beaulieu, J.M. 2021). Likelihood-based methods utilise additional information from branch lengths, which is frequently cited as a reason to prefer these methods over parsimony-based approaches (Schluter, D., Price, T., et al. 1997, Royer-Carenzi, M., Pontarotti, P., et al. 2013, Wright, A.M. and Hillis, D.M. 2014, Joy, J.B., Liang, R.H., et al. 2016). However, this argument neglects the fact that likelihood methods require estimation of additional parameters –the transition rates of an evolutionary model– which are not necessary in a parsimony approach. More complex models will require more transition rates to be estimated from the data, thereby spreading the available information increasingly thinly (Mooers, A.Ø. and Schluter, D. 1999). The difficulty in co-estimating both ancestral states and transition rates has been highlighted in a recent study by Gascuel, O. and Steel, M. (2020). They show that evolutionary histories which provide optimal information for estimating pattern (i.e. ancestral states) vs. process (i.e. transition rates) are mutually exclusive. Accuracy of ancestral state estimates is further confounded by a number of interrelated tree properties: e.g. size, shape, ultrametricity, branch length distribution (Mooers, A.Ø. 2004, Li, G., Steel, M., et al. 2008, Royer-Carenzi, M., Pontarotti, P., et al. 2013). These properties may reflect evolutionary history but can equally reflect taxon selection bias. For example, inclusion/exclusion of fossil tips can affect all aforementioned tree properties.

Estimating accurate ancestral states from discrete data requires selecting an appropriate method/model. Model selection is standard practice in likelihood-based ancestral state estimation, and is typically achieved by comparing log likelihood scores using information criteria such as the Akaike Information Criterion (AIC, Akaike, H. (1973)) or Bayesian Information Criterion (BIC, Schwarz, G. (1978)). Previous simulation studies have demonstrated the effectiveness of these methods in discriminating between competing models of sequence evolution (Posada, D. and Crandall, K.A. 2001, Jhwueng, D.-C., Huzurbazar, S., et al. 2014). However, their efficacy in the context of ancestral state estimation has never been explicitly tested. Information criteria are sensitive to sample size (Posada, D. and Buckley, T.R. 2004, Dziak, J.J., Coffman, D.L., et al. 2020, Susko, E. and Roger, A.J. 2020). Thus, the paucity of data used to estimate ancestral states (i.e. a single character compared with hundreds or thousands of sites) may prohibit accurate model selection, particularly for small trees with few tips. There is no statistical method for selecting between likelihood and parsimony approaches and consequently a simulation approach is required to compare these methods. Previous simulation studies suggest that likelihood-based ancestral state estimation tends to yield more accurate results than parsimony (Gascuel, O. and Steel, M. 2014, Holland, B.R., Ketelaar-Jones, S., et al. 2020). It is important to note, however, that these studies have primarily focused on large ultrametric trees with over 100 tips. As a result, the extent to which these findings can be generalised remains uncertain.

Evidently, a better understanding of the relative merits and limitations of different methods of ancestral state estimation is overdue. In this study, I employ a simulation-based approach to compare the performance of parsimony and likelihood methods in estimating ancestral states from discrete binary characters. To achieve this, I simulate characters using 15 evolutionary models encompassing a range of transition rates (including both slow to fast and equal to asymmetrical rates). I test tree sizes ranging from 8 to 256 tips, and three different topologies representing extremes of tree symmetry and branch length heterogeneity. Using the simulated tip states, I estimate ancestral states under parsimony assuming accelerated transformations (ACCTRAN), and under maximum likelihood using the Equal Rates (ER) and All-Rates-Different (ARD) models implemented using several optimisation methods. In doing so I test the following: 1) How does the number of tips affect the accuracy of different methods? 2) How does tree shape affect the accuracy of different methods? 3) How does branch length heterogeneity affect the accuracy of different methods? 4) How does evolutionary transition rate affect the accuracy of different methods? 5) How does evolutionary transition rate asymmetry affect the accuracy of different methods? 6) How does the optimisation method affect the accuracy of likelihood-based models? Additionally, I compare the difference in accuracy between ER and ARD models with the corresponding differences in model likelihood and model uncertainty to test the following: 7) Does model likelihood predict accuracy? 8) Does model uncertainty predict accuracy? Finally, I compare parsimony and likelihood ancestral state estimation of four published morphological character matrices, evaluating the results in light of the simulation experiments. The results of this study provide valuable insights into the accuracy of parsimony and maximum likelihood methods for estimating ancestral states, thereby informing best practice for future evolutionary studies.

## Methods

### Data simulation

I simulated data using trees of 8, 16, 32, 64, 128, and 256 tips. These reflect tree sizes that are common for morphological datasets (O’Leary, M. and Kaufman, S. 2012). Data were simulated using three topologies: fully symmetrical (balanced) ultrametric tree with equal branch lengths; fully asymmetrical (unbalanced) tree with equal branch lengths and fully asymmetrical ultrametric tree with equal internal branch lengths. These topologies represent end members both in terms of tree symmetry and branch length heterogeneity. All trees were fixed to a constant height of 1. For each topology and tree size, I iterated over five distinct rate categories (0.2, 0.4, 0.6, 0.8, 1) to create fifteen discrete Markov models with varying forward and backward transition rates. All of the resulting Markov models fall within three end members: slow equal rates (instantaneous rates = 0.2); fast equal rates (instantaneous rates = 1); highly unequal rates (instantaneous rates = 0.2 & 1). I conducted 270 discrete experiments (i.e. unique combinations of tree size, topology and Markov model). In each, I simulated replicate binary characters, discarding parsimony uninformative characters (i.e. characters with states that were not represented by at least two tips) until a total of 500 characters was reached. This equates to a total of 135,000 simulated characters over all experiments. Data simulation was conducted using the R programming language (R Core Team 2023). Fully symmetrical and asymmetrical trees were created using the read.newick() function from the ‘phytools’ package (Revell, L.J. 2012). Binary characters were simulated using the simulate_mk_model() function from the ‘castor’ package (Louca, S. and Doebeli, M. 2018), specifying flat (equal) root probabilities.

### Ancestral state estimation

Simulated tip states were then subjected to ancestral state estimation. Ancestral states were estimated under parsimony with accelerated transformations (ACCTRAN) using the function ancPropStateMat() from the ‘paleotree’ package (Bapst, D.W. 2012), which is a wrapper for the ancestral.pars() function from the ‘phangorn’ package (Schliep, K.P. 2011). Likelihood ancestral state estimation was conducted primarily using the corHMM() function from the ‘corHMM’ package (Boyko, J.D. and Beaulieu, J.M. 2021). I used the Equal Rates (ER) and All Rates Different (ARD) models with one rate category, root probabilities estimated using the method of Yang, Z. (2006) and random restarts set to 10. I also estimated ancestral states using the true model (i.e. the model under which the data were simulated). The corHMM() function utilises marginal likelihood estimation, yet alternative likelihood optimisation methods are frequently applied when estimating ancestral states. To compare likelihood optimisation methods, I repeated the ER and ARD analyses using stochastic mapping implemented in ‘phytools’ (Revell, L.J. 2012) and scaled conditional likelihood estimation implemented in the ‘ape’ R package (Paradis, E. and Schliep, K. 2019). All ancestral states were estimated using the true tree (i.e. the tree on which the data were simulated).

I selected the best fitting model between ER and ARD models (optimised using corHMM) using AIC and BIC comparison. I also calculated the difference in model uncertainty between ER and ARD models by comparing raw uncertainty scores (see measuring error and uncertainty below).

### Measuring error and uncertainty

Ancestral state estimation methods typically output a matrix, ***A***, of dimensions ***S*** by ***N***, where ***S*** denotes the number of observed states (=2 in all simulations presented here) and ***N*** denotes the number of nodes present in the tree used for estimation. Each cell contains a value between 0 and 1 corresponding to the probability that a given state was ancestrally present at a given node. States are mutually exclusive, thus rows sum to 1. Under marginal likelihood estimation, states can be assigned any probability. In contrast, parsimony reconstructs states as either unambiguously present; unambiguously absent, or equally uncertain between two or more states. For binary characters, states are thus reconstructed as either present (probability = 1); absent (probability = 0) or ambiguous (probability = 0.5).

I measured the error of ancestral state estimates using two strictly proper scoring metrics. The first is ‘raw error’ (Holland, B.R., Ketelaar-Jones, S., et al. 2020). Raw error is a simple linear scoring and is calculated by summing the probabilities assigned to incorrect states (which is equal to one minus the probability assigned to the correct state) for each node, and then dividing this sum by the total number of nodes:

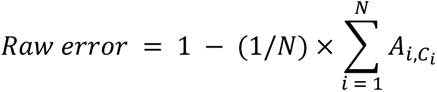

Where ***C*** is a vector of correct states of length ***N***.

Under this scoring, a completely accurate estimate would receive a score of zero, whereas a completely inaccurate estimate would receive a score of one. I also calculated Brier score (Brier, G.W. 1950), a popular metric for evaluating the error of probabilistic estimates, which is defined as the mean squared difference between probabilistic estimates assigned to a set of events and the corresponding true outcomes of those events. Lower Brier scores indicate better estimates: for binary data a score of zero denotes perfect accuracy, whereas a score of one denotes a completely incorrect estimate. Compared to the raw score measure, Brier scoring less strongly penalises uncertainty (see Supplementary Fig. 1). I calculated uncertainty of ancestral state estimates using equivalent measures: “Raw uncertainty” is a simple linear measure calculated by summing the highest probabilities assigned to any state for each node, dividing this by the total number of nodes and then subtracting this value from one:

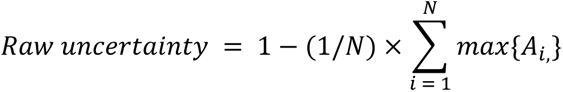

A score of zero indicates a completely certain ancestral state estimate (i.e. one state is assigned maximum probability for each node). Conversely, a score of one represents a completely uncertain ancestral state estimate (i.e. all states are equally probable for each node). I also calculated Shannon entropy (Shannon, C.E. 1948), which measures the average information, or ‘surprise’, in a probability distribution. For a comparison between raw uncertainty and entropy, see Supplementary Fig. 1. Brier score and entropy were calculated using the ‘measures’ and ‘DescTools’ R packages respectively (Signorell, A., Aho, K., et al. 2019, Probst, P. 2022). Finally, for ER and ARD likelihood models, I calculated the sum of estimated transition rates.

### Comparing model selection criteria

I compared the performance of three model selection criteria: difference in AIC score (ΔAIC), difference in BIC score (ΔBIC) and difference in raw uncertainty (Δraw uncertainty). I used statistical classification theory to quantify the diagnostic power of these criteria in rejecting the ER model in favour of the more highly parameterized ARD model. I computed a confusion matrix for each criterion, classifying each result as a false positive/type II error (ARD selected, ARD has higher raw error); false negative/type I error (ARD rejected, ER has higher raw error); true positive (ARD selected, ER has higher raw error); true negative (ARD rejected, ARD has higher raw error). I then calculated the test sensitivity (or true positive rate), defined as the number of true positives divided by the sum of true positives + false positives; test specificity (or true negative rate), defined as the number of true negatives divided by the sum of true negatives + false negatives; and test efficiency (or diagnostic effectiveness), defined as the sum of true positives and true negatives divided by the sum of true positives, true negatives, false positives and false negatives (Šimundić, A.-M. 2009).

### Empirical study

I evaluated the effect of different ancestral state estimation methods on empirical morphological data. I selected four published morphological character-by-taxon matrices with associated time scaled phylogenies. These matrices cover early gnathostomes (King, B., Qiao, T., et al. 2017); early arthropods (Wolfe, J.M. 2017); mammals (Lee, M.S. 2016) and theropod dinosaurs (Bapst, D., Wright, A., et al. 2016). Unlike simulated characters, empirical matrices contain missing data (i.e. character states that haven’t been observed in a particular taxon). Taxa may also be coded as uncertain or polymorphic between two or more states. In corHMM, such ambiguities can be accommodated by coding possible states separated by the ‘&’ symbol. The function then assigns likelihoods to possible states following Felsenstein, J. (2004 pg. 255). For parsimony analyses, uncertain tip states were assigned possible states using a contrast matrix via the ‘contrast’ argument of the ancPropStateMat() function from paleotree (Bapst, D.W. 2012). I subsampled characters from each matrix, selecting only those that were coded for >= 32 tips, thus mitigating biases associated with characters containing very high amounts of missing data.

It is impossible to determine the error of an empirical ancestral state estimate given that the true evolutionary history is unknown. Instead, I compared the uncertainty of ancestral state estimates between methods. I also measured incongruence between ancestral state estimation methods using the following approaches: Firstly, I considered the difference in root state estimates. Methods were considered to have estimated different root states if both estimates were certain (<= ⅓ uncertainty) and assigned maximum probability to different states. I subsampled characters from each taxon-by-character matrix, retaining only characters for which the root state had been estimated with low uncertainty (<= ⅓) across all estimation methods. By comparing the most likely root state of each estimation method, I was able to classify each character into one of four mutually exclusive categories: 1) all methods infer the same root state; 2) parsimony infers a different root state to ER and ARD; 3) ER infers a different root state to parsimony and ARD; 4) ARD infers a different root state to parsimony and ER. Secondly, I considered the difference between estimation methods across all ancestral states (including the root state). Methods were considered to have estimated different ancestral states if both methods shared at least a single node with low uncertainty (<= ⅓ uncertainty) but assigned maximum probability to different states for that node. To achieve this, I performed a subsampling of characters from each taxon-by-character matrix, selecting only those characters for which at least one node was estimated with low uncertainty (<= ⅓) across all estimation methods. I then classified each character into one of five mutually exclusive categories: 1) all methods infer compatible evolutionary histories; 2) all methods infer incompatible evolutionary histories; 3) parsimony infers a different evolutionary history to ER and ARD; 4) ER infers a different evolutionary history to parsimony and ARD; 5) ARD infers a different evolutionary history to parsimony and ER. All characters were treated as unordered.

## Results

Relative and absolute ancestral state estimation error varies with method, tree size, tree shape, branch length heterogeneity and transition rates of the simulating model (Fig. 1, Supplementary Figs. 2, 3). The ER model performs best overall; however, it does not perform optimally in every experiment. Furthermore, the relative performance of different methods depends on the measure of error used to compare them (Supplementary Figs. 2, 3). This reflects differences in the uncertainty of ancestral estimates between methods. Parsimony typically exhibits the lowest ancestral state estimation uncertainty (Fig. 2, Supplementary Figs. 4, 5), which is unsurprising given how parsimony optimises ancestral states. The ARD model typically exhibits the highest uncertainty (Fig. 2, Supplementary Figs. 4, 5). Absolute uncertainty varies with the aforementioned data properties, although relative uncertainty remains largely consistent across all experiments (Fig. 2, Supplementary Figs. 4, 5). Comparing the likelihood models, the ARD model tends to show higher error than the ER model. ARD exhibits higher mean raw error in 225/270 experiments and higher mean Brier score in 182/270 experiments (Supplementary Figs. 2, 3). Estimated transition rates vary between likelihood models, and covary with tree shape and size. The ARD model tends to estimate higher transition rates than the ER model, thus incurring greater transition rate error (Supplementary Fig. 6).

**Figure 1.**
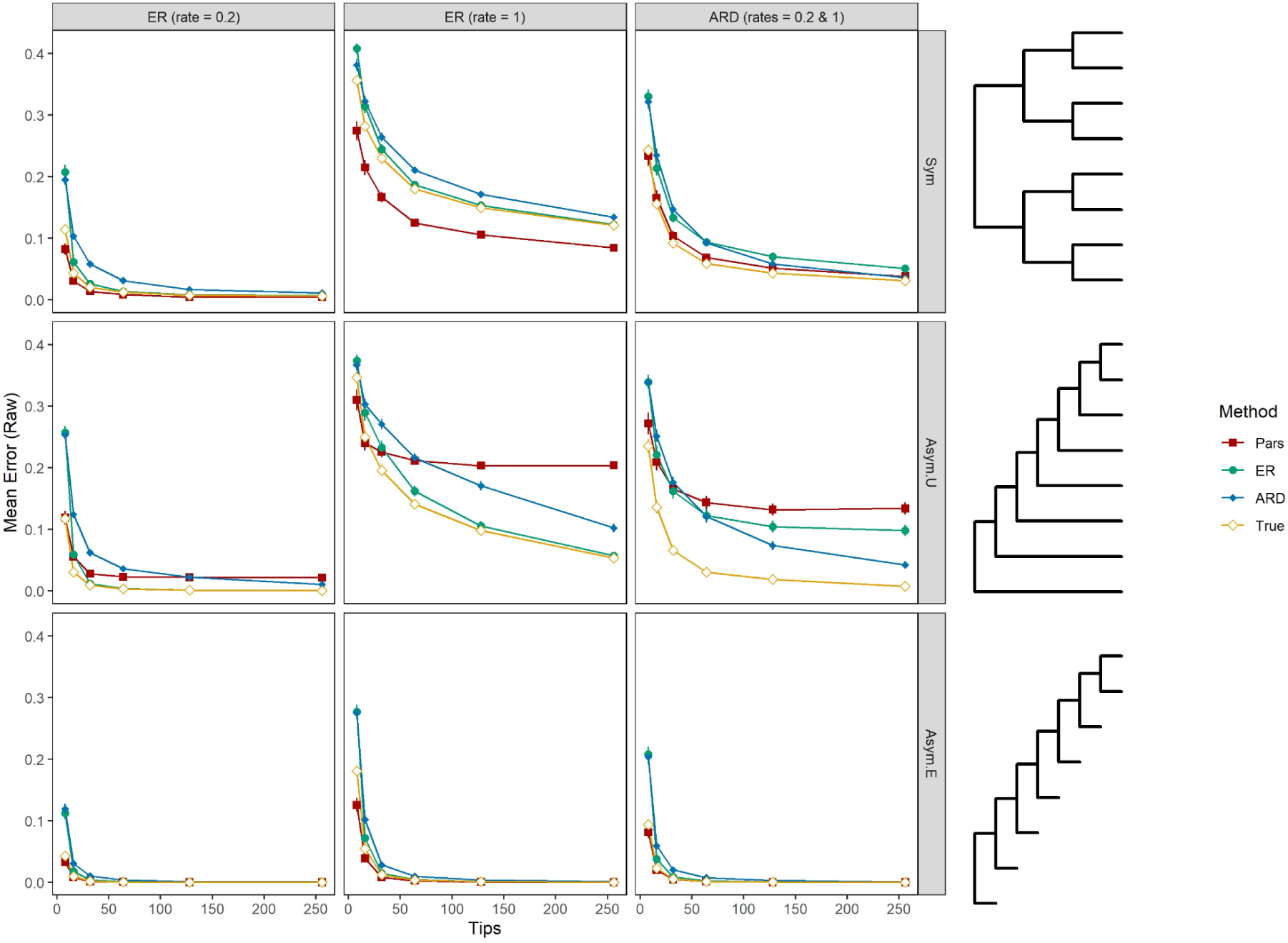
Mean raw error of different ancestral state estimation methods plotted against the number of tips present in the tree used to simulate character data: ACCTRAN Parsimony (Pars); Equal Rates model (ER); All-Rates-Different model (ARD); the true model under which the data were simulated (True). Results are presented for data generated under three different models: Equal Rates (rate = 0.2); Equal Rates (rate = 1); All-Rates-Different (rates = 0.2 & 1) and three different tree shapes: symmetrical equal branch lengths (Sym); asymmetrical ultrametric (Asym.U); and asymmetrical equal branch lengths (Asym.E). Each data point represents the mean error of 500 simulated characters. Error bars show the 95% confidence interval of the mean. All methods exhibit less error when more tips are present. Parsimony has low mean error when branch lengths are equal, but is outperformed by likelihood models when branch lengths are unequal.

**Figure 2.**
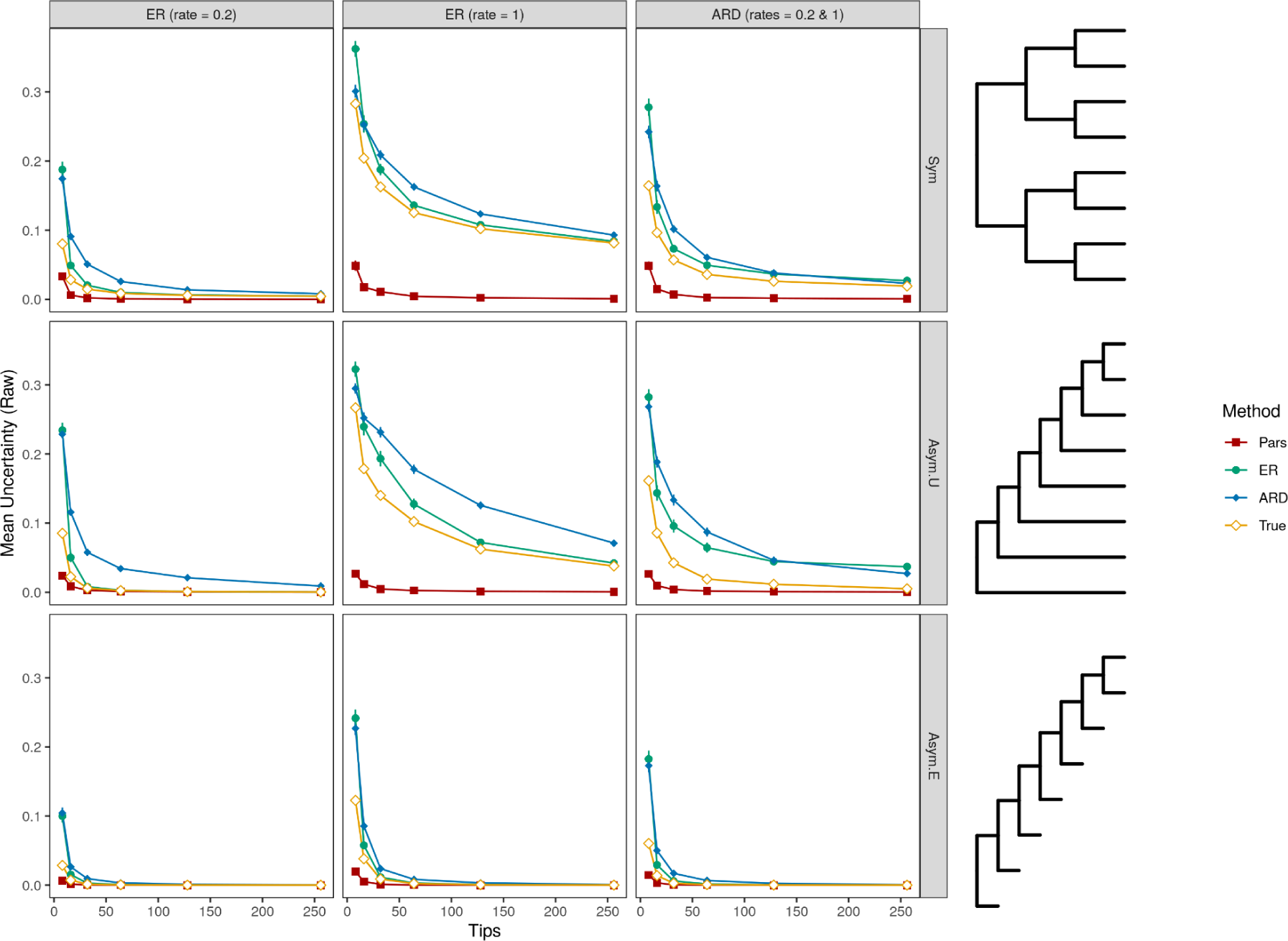
Mean raw uncertainty of different ancestral state estimation methods plotted against the number of tips present in the tree used to simulate character data: ACCTRAN Parsimony (Pars); Equal Rates model (ER); All-Rates-Different model (ARD); the true model under which the data were simulated (True). Results are presented for data generated under three different models: Equal Rates (rate = 0.2); Equal Rates (rate = 1); All-Rates-Different (rates = 0.2 & 1) and three different tree shapes: symmetrical equal branch lengths (Sym); asymmetrical ultrametric (Asym.U); and asymmetrical equal branch lengths (Asym.E). Each data point represents the mean error of 500 simulated characters. Error bars show the 95% confidence interval of the mean. All methods exhibit less uncertainty when more tips are present. ARD typically has slightly higher mean uncertainty, while parsimony exhibits much lower mean uncertainty than all other methods.

### Effect of tree size

Tree size (i.e. the number of tips) is negatively correlated with mean ancestral state estimation error, calculated using either the raw or Brier scores. The relationship is nonlinear; error drops rapidly if tips are added to small trees (less than ≈64 tips). Error reduces more gradually as tree size increases (Fig. 1, Supplementary Figs. 2, 3). A similar relationship is observed between tree size and mean uncertainty, calculated using either the raw or entropy scores (Fig. 2, Supplementary Figs. 4, 5). Tree size is also negatively correlated with the mean sum of estimated transition rates (Supplementary Fig. 6).

### Effect of tree shape and branch length heterogeneity

All methods exhibit lower error when estimating ancestral states using the non-ultrametric tree (i.e. the asymmetrical tree with equal branch lengths, Fig 1, Supplementary Figs. 2, 3, 7, 8). Parsimony exhibits high error when applied to the asymmetrical ultrametric tree and is outperformed by both likelihood models as the number of tips increases (Fig 1. Supplementary Figs. 2, 3, 7, 8). Likelihood models show slightly higher error when applied to symmetrical vs. asymmetrical ultrametric trees (Supplementary Figs. 7, 8). Parsimony exhibits low uncertainty regardless of tree shape or branch length heterogeneity (Supplementary Figs. 9, 10). Likelihood models show similar distributions of uncertainty when applied to symmetrical vs. asymmetrical ultrametric trees (Supplementary Figs. 9, 10).

### Effect of transition rates

All methods show a positive correlation between the sum of transition rates used to generate the data and the mean error of the ancestral state estimates measured using either the raw or Brier score: Ancestral state estimation methods exhibit more error when applied to data generated under faster transition rates (Fig 3, Supplementary Fig. 11). The relationship between transition rate heterogeneity and method estimation error is less obvious. For some experiments, particularly those conducted using the asymmetrical ultrametric tree, parsimony and ER methods show a positive correlation between transition rate heterogeneity and estimation error measured using either the raw or Brier score (Fig 3, Supplementary Fig. 12). This suggests that as transition rate heterogeneity increases parsimony and ER ancestral state estimates are more prone to error. In contrast, the error of ARD estimates remains fairly constant as transition rate heterogeneity increases (Fig 3, Supplementary Fig. 12). With the exception of parsimony, the uncertainty of ancestral state estimation methods is positively correlated with the sum of transition rates used to generate the data: likelihood ancestral state estimation methods exhibit more uncertainty when applied to data generated under faster transition rates (Supplementary Figs. 13, 14). The ARD model shows a negative correlation between uncertainty and transition rate heterogeneity: uncertainty decreases as transition rate heterogeneity increases. In comparison, ER shows a negative correlation when applied to the symmetrical ultrametric tree but shows a positive correlation when applied to the asymmetrical ultrametric tree (Supplementary Figs. 15, 16).

**Figure 3.**
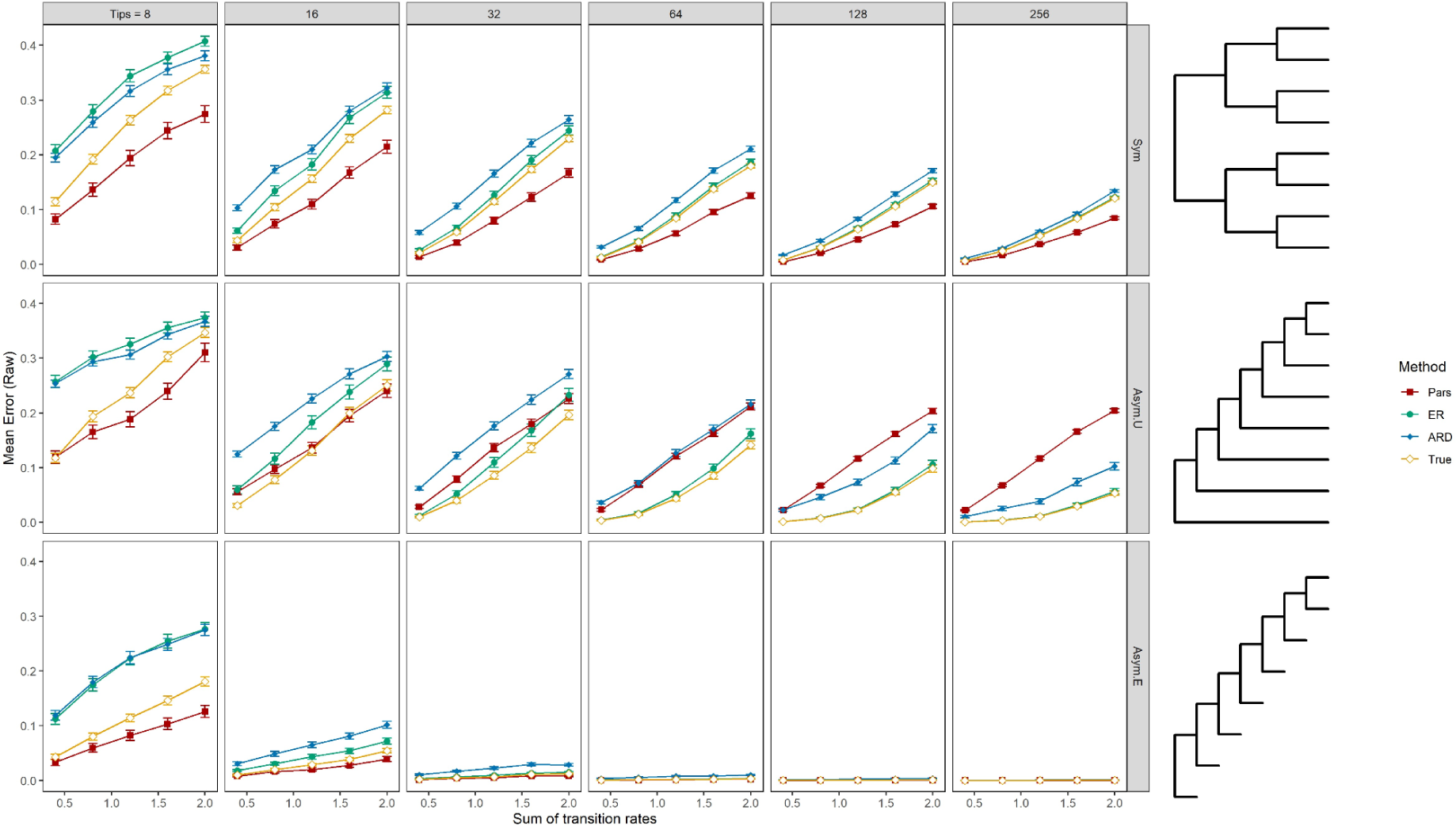
Mean raw error of different ancestral states estimation methods plotted against the sum of transition rates used to simulate character data: ACCTRAN Parsimony (Pars); Equal Rates model (ER); All-Rates-Different model (ARD); the true model under which the data were simulated (True). Each data point represents the mean of 500 binary characters generated under an ER model. Error bars show the 95% confidence interval of the mean. Results are plotted for trees of size 8-256 tips using three topologies: symmetrical equal branch lengths (Sym); asymmetrical ultrametric (Asym.U); asymmetrical equal branch lengths (Asym.E). All methods exhibit less error when the sum of transition rates is lower.

**Figure 4.**
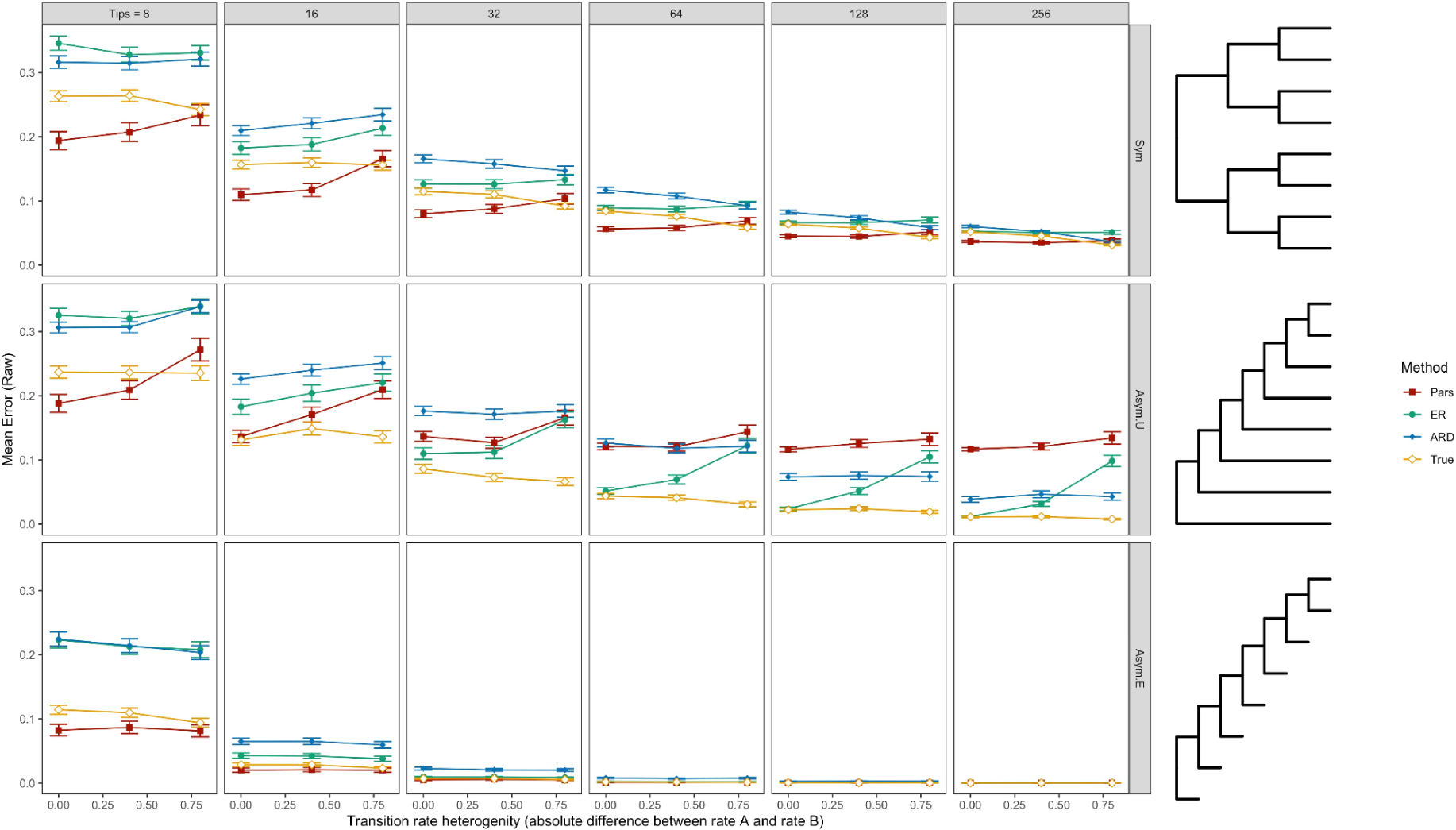
Mean raw error of different ancestral states estimation methods plotted against the transition rate heterogeneity used to simulate character data, calculated by taking the absolute difference between forward and backward transition rates. ACCTRAN Parsimony (Pars); Equal Rates model (ER); All-Rates-Different model (ARD); the true model under which the data were simulated (True). Each data point represents the mean of 500 binary characters generated under a model in which the sum of transition rates is equal to 1.2. Error bars show the 95% confidence interval of the mean. Results are plotted for trees of size 8-256 tips using three topologies: symmetrical equal branch lengths (Sym); asymmetrical ultrametric (Asym.U); asymmetrical equal branch lengths (Asym.E). Under ultrametric trees, ER and parsimony error tends to marginally increase as transition rate heterogeneity increases.

### Effect of likelihood optimisation

Different optimisation methods exhibit virtually identical error (measured using either raw or Brier score) when using the ER model (Sup Fig. 17, 18). However, error varies with likelihood optimisation method when using the ARD model (Fig 5, Sup Figs. 19, 20). Marginal likelihood estimation using the ‘corHMM’ package is consistently less erroneous than either stochastic mapping using ‘phytools’ or conditional likelihood estimation using ‘ape’. Conditional likelihood estimation tends to exhibit the highest error. Difference in error between optimisation methods is greatest when applied to asymmetrical ultrametric trees (Fig 5, Sup Fig. 19, 20).

**Figure 5.**
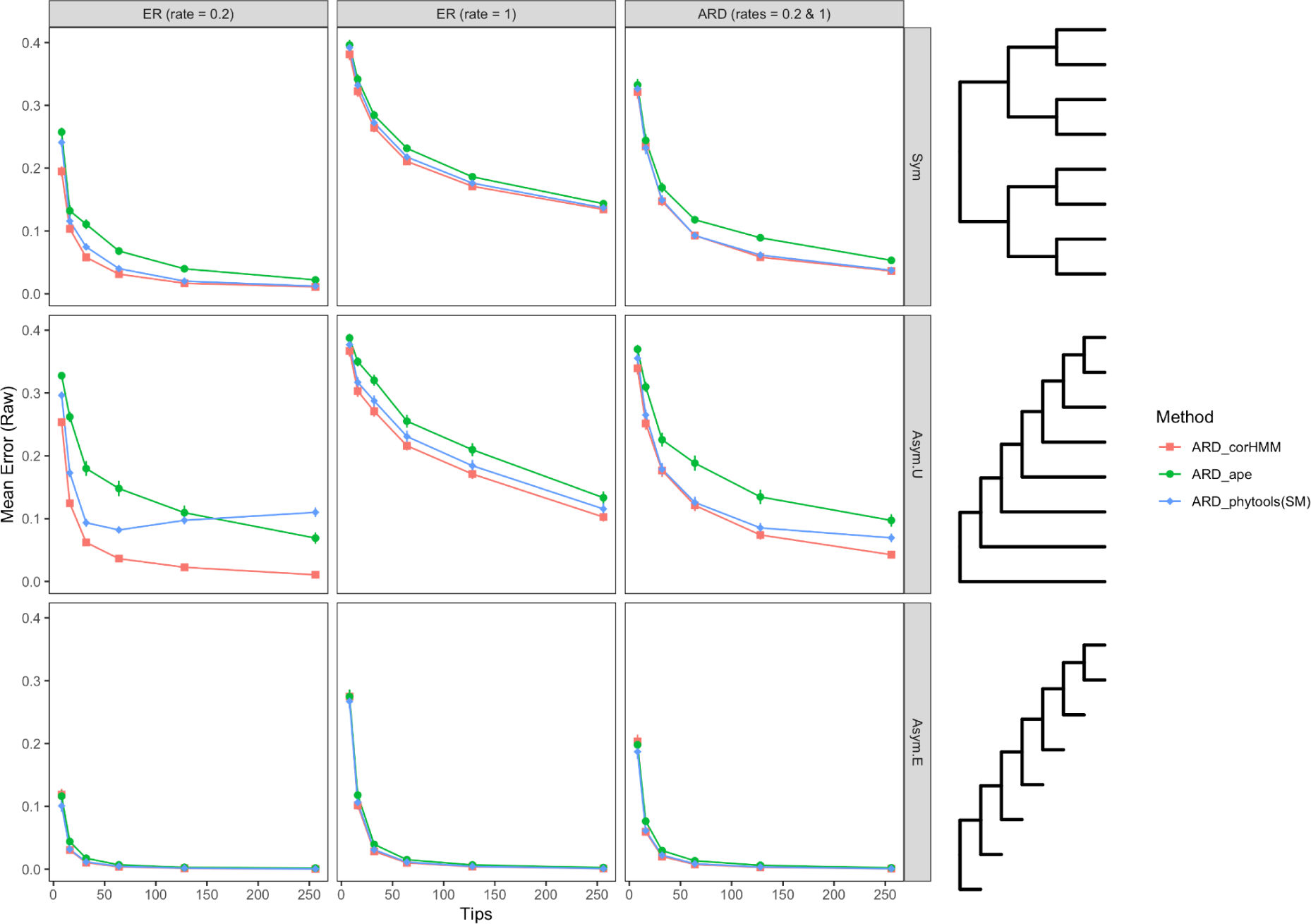
Mean raw error of the ARD model optimised using marginal likelihood (corHMM, red), scaled conditional likelihood (ape, green) and stochastic mapping (phytools, blue) plotted against the number of tips present in the tree used to simulate character data. Results are presented for data generated under three different models: Equal Rates (rate = 0.2); Equal Rates (rate = 1); All-Rates-Different (rates = 0.2 & 1) and three different tree shapes: symmetrical equal branch lengths (Sym); asymmetrical ultrametric (Asym.U); and asymmetrical equal branch lengths (Asym.E). Each data point represents the mean error of 500 simulated characters. Error bars show the 95% confidence interval of the mean. Marginal likelihood optimisation implemented in corHMM consistently performs better than alternative optimisation methods.

### Performance of model selection criteria

Difference in raw uncertainty between ER and ARD models (Δraw uncertainty) is strongly positively correlated with difference in raw error (Fig. 6). A model with higher uncertainty tends to estimate ancestral states with higher error. The greater the difference in uncertainty between models, the greater the difference in error. These measures are somewhat autocorrelated; very uncertain estimates will inevitably exhibit more error. Yet the observed correlation is stronger than would be expected due to autocorrelation alone (Fig. 6). The relationship is a zero intercept correlation, meaning that the line of best fit passes through the origin. Note that raw uncertainty is not correlated with raw error under parsimony ancestral state estimation, which universally exhibits low raw uncertainty irrespective of error (Supplementary Fig. 9). Difference in AIC score between ER and ARD models (ΔAIC) is weakly negatively correlated with difference in raw error, with a non zero intercept (Fig. 7). This result contradicts the expectation that models with a better (lower) AIC score will be more accurate (and vice versa), instead suggesting the opposite will occur in some instances. Using ΔBIC, rather than ΔAIC, results in shifting the distribution to the left, as the simpler ER model is selected more frequently (Supplementary Fig. 21). Comparing the diagnostic power of these criteria, Δraw uncertainty has higher test sensitivity, specificity and efficiency than both ΔAIC and ΔBIC (Table 1). In other words, difference in raw uncertainty is a better predictor of model accuracy than difference in AIC or BIC.

**Figure 6.**
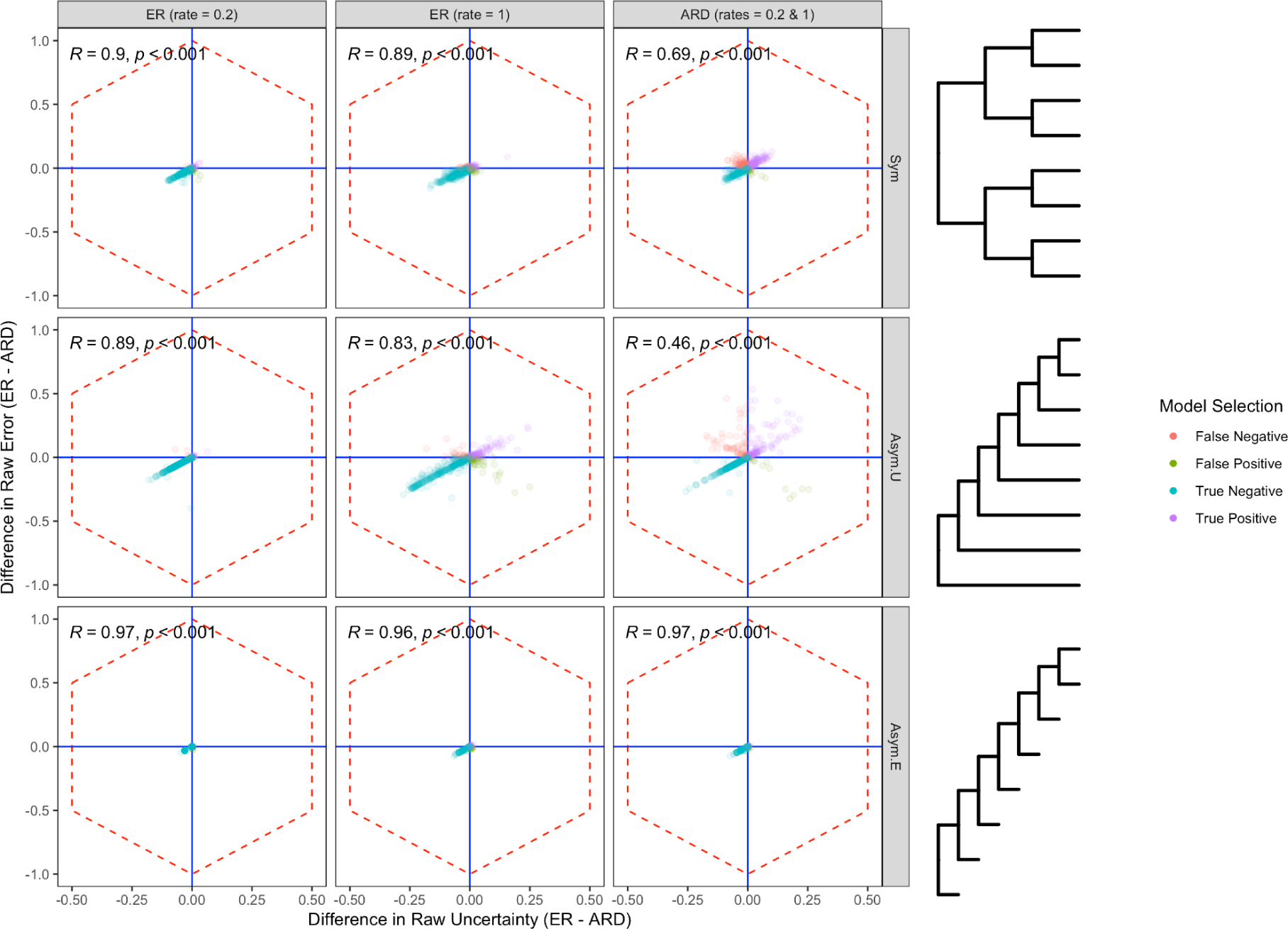
Difference in raw uncertainty between ER and ARD models (Δraw uncertainty) plotted against the difference in raw error between ER and ARD models (Δraw error). Results are presented for data generated under three different models: Equal Rates (rate = 0.2); Equal Rates (rate = 1); All-Rates-Different (rates = 0.2 & 1) using 64-tip trees of three different shapes: symmetrical equal branch lengths (Sym); asymmetrical ultrametric (Asym.U); and asymmetrical equal branch lengths (Asym.E). Top = ER has more error; Bottom = ARD has more error; Left = ARD has more uncertainty; Right = ER has more uncertainty. Hexagons mark limits of possible plot space due to autocorrelation.

**Figure 7.**
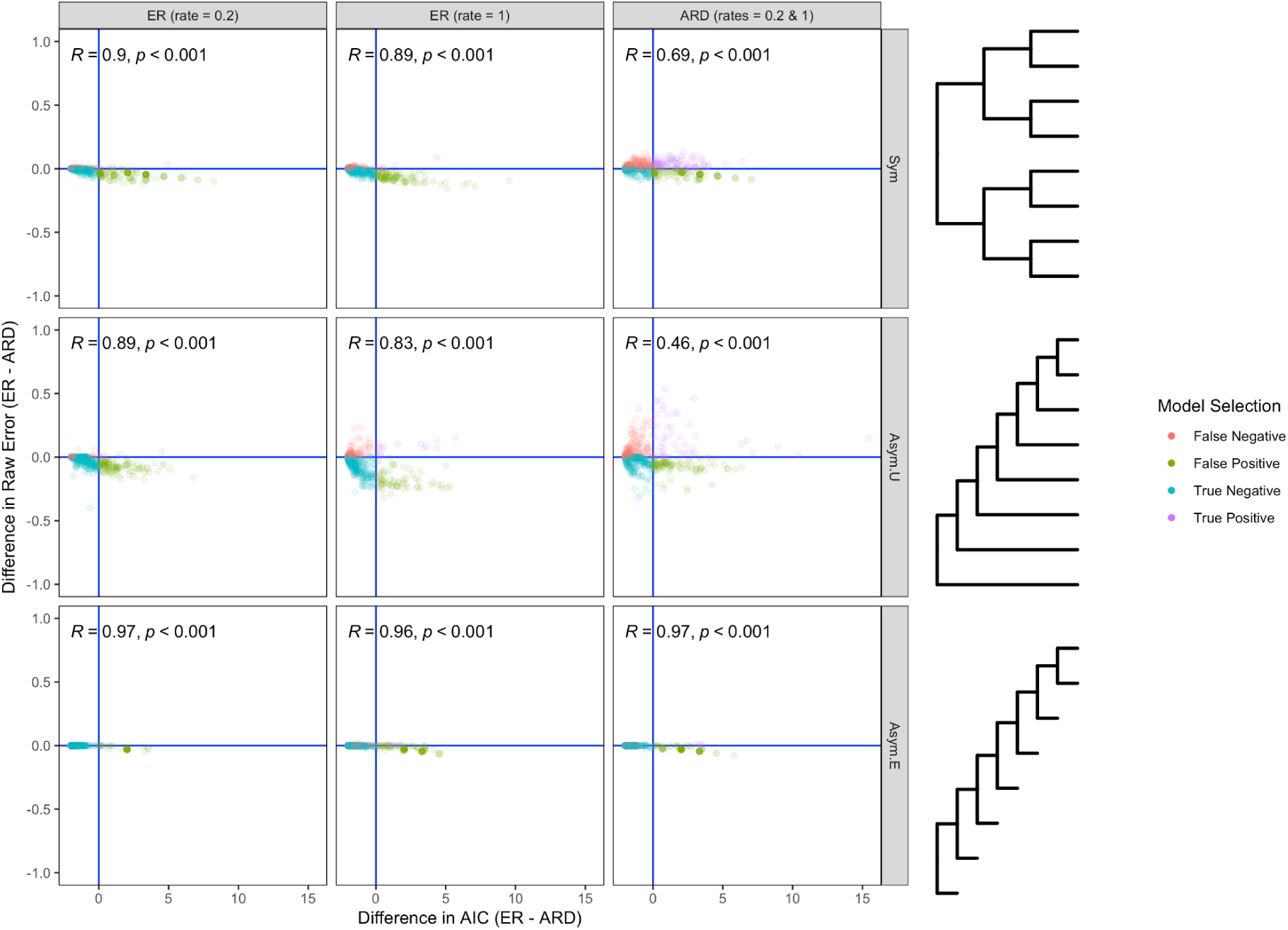
Difference in AIC score between ER and ARD models (ΔAIC) plotted against the difference in raw error between ER and ARD models (Δraw error). Results are presented for data generated under three different models: Equal Rates (rate = 0.2); Equal Rates (rate = 1); All-Rates-Different (rates = 0.2 & 1) using 64-tip trees of three different shapes: symmetrical equal branch lengths (Sym); asymmetrical ultrametric (Asym.U); and asymmetrical equal branch lengths (Asym.E). Top = ER has more error; Bottom = ARD has more error; Left = ER has better AIC score; Right = ARD has better AIC score.

**Table 1.** Comparison of the diagnostic power of ΔAIC, ΔBIC and Δraw uncertainty for rejecting the ER model in favour of the ARD model. Results are presented for 135000 analyses (270 experiments * 500 replicates). Note that in 241 analyses, ER and ARD models exhibited identical raw error. These results are thus excluded.

| | $\Delta AIC$ | $\Delta BIC$ | $\Delta \text{Raw Uncertainty}$ |
| --- | --- | --- | --- |
| False Negative (Type II error) | 31680 | 35007 | 7398 |
| False Positive (Type I error) | 21602 | 8342 | 8195 |
| True Negative | 74379 | 87639 | 87786 |
| True Positive | 7098 | 3771 | 31380 |
| Sensitivity | 0.25 | 0.31 | 0.79 |
| Specificity | 0.70 | 0.71 | 0.92 |
| Efficiency | 0.60 | 0.68 | 0.88 |

### Empirical study

Comparison of ancestral state estimation methods using empirical character matrices reveals consistent differences in uncertainty and incongruence between methods (Fig. 8). Parsimony always exhibits the lowest uncertainty. ARD exhibits a slightly higher average raw uncertainty than ER in three of the four datasets (Fig. 7E-H). This is consistent with the results from the simulated data (Fig. 2, Supplementary Figs. 4). Across all datasets, root state estimates are largely identical, regardless of estimation method (congruence: Arthropods = 96.5%, Dinosaurs = 99.6%, Gnathostomes = 100%, Mammals = 98%). Rare differences are due to parsimony or ARD estimating a unique root state (Fig. 7I-L). Considering all ancestral states rather than just the root state, methods are more frequently incongruent (congruence: Arthropods = 66.7%, Dinosaurs = 61%, Gnathostomes = 64.6%, Mammals = 63%). The most frequent cause of this incongruence is that parsimony ancestral state estimates are incompatible with ER and ARD estimates. The second most frequent cause of incongruence is that ARD estimates are incompatible with those of parsimony and ER (Fig. 7M-P). Taken together, these results show that, when applied to empirical data, different ancestral state estimation methods yield substantively different results, both in terms of their precision and inferred evolutionary history.

**Figure 8.**
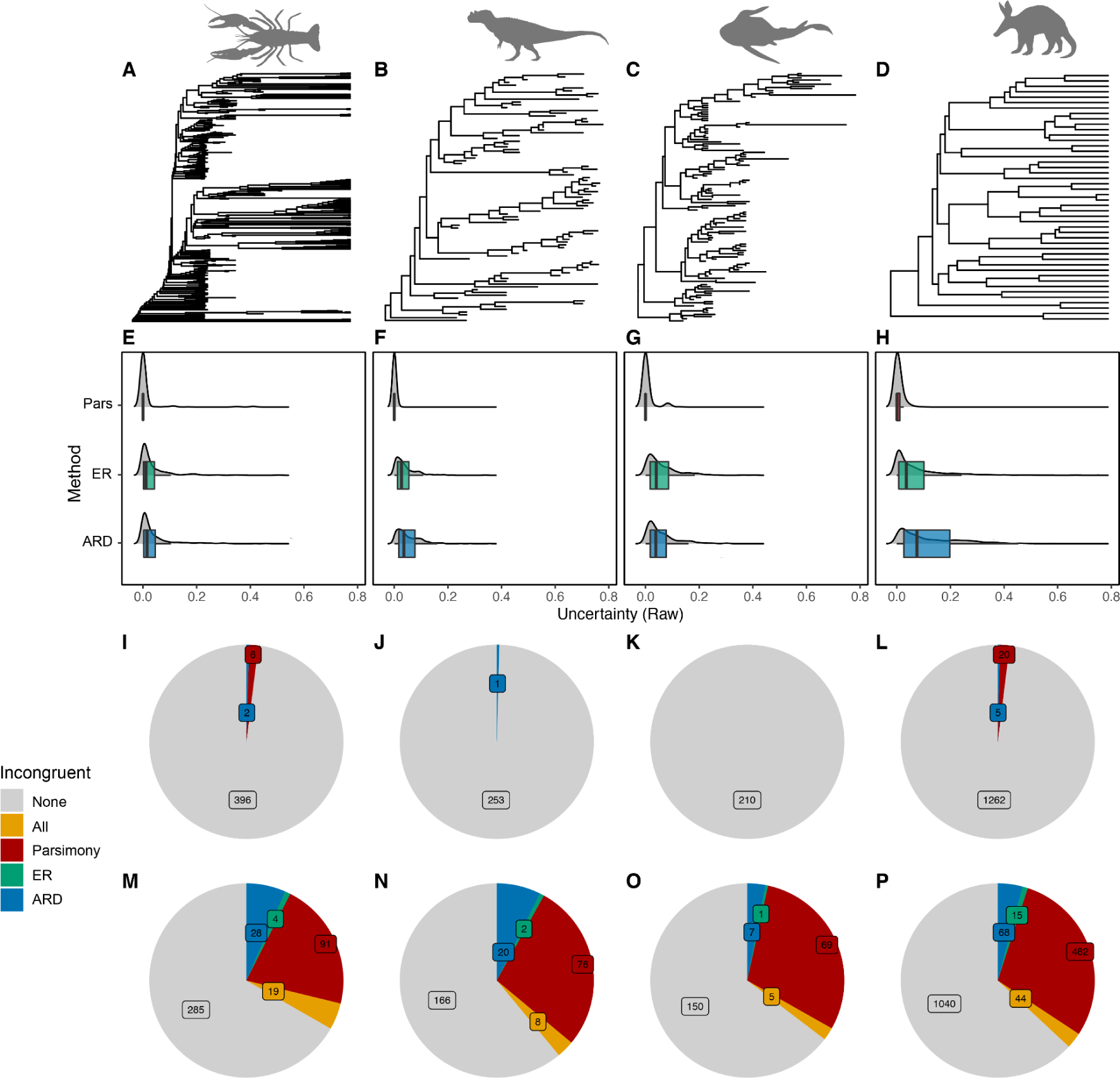
Ancestral state estimation results for four empirical morphological datasets: A) Arthropods (Wolfe, J.M. 2017); B) dinosaurs (Bapst, D., Wright, A., et al. 2016); C) Early gnathostomes (King, B., Qiao, T., et al. 2017); Mammals (Lee, M.S. 2016). E-H) Uncertainty of ancestral state estimates using Parsimony (Pars), Equal Rates (ER) and All Rates Different (ARD). I-L) Pie charts showing frequency of incongruence among root state estimates: Grey = all methods infer the same root state; Red = parsimony infers a different root state to ER and ARD; Green = ER infers a different root state to parsimony and ARD; Blue = ARD infers a different root state to parsimony and ER. M-P) Pie charts showing frequency of incongruence among all ancestral state estimates: Grey = all methods infer compatible evolutionary histories; Yellow = all methods infer incompatible evolutionary histories; Red = parsimony infers a different evolutionary history to ER and ARD; Green = ER infers a different evolutionary history to parsimony and ARD; Blue = ARD infers a different evolutionary history to parsimony and ER.

## Discussion

### Ancestral state estimation using the Equal Rates model is most accurate most of the time

The ER model performs consistently well under most simulated scenarios, even with small trees and when the simulated evolutionary process violates the assumption of equal rates (Supplementary Figs. 2,3). The ER model also rarely infers evolutionary histories from empirical characters that are unique (i.e. at odds with either parsimony or ARD ancestral state estimates, Fig. 8). ER ancestral state estimates are typically supported by alternative methods. These results suggest that the ER model provides a robust benchmark against which ancestral estimates derived using alternative methods should be critically considered. That said, the relative and absolute performance of all methods depends a great deal on data attributes related to both sampling and the evolutionary process that generated the data. To further complicate matters, relative performance of different methods is subjective as it depends on the scoring metric used (or more specifically, how uncertainty is penalised by different scoring metrics). This can be demonstrated with a contrived example: Consider a binary character evolving on a tree with four internal nodes. The ancestral states of these nodes have been estimated using two methods. Method A estimates that each node has a probability of 0.8 of being state 1. Method B estimates three of the nodes as state 1, with a probability of 1, with the remaining node estimated as ambiguous between state 1 and 2 (probability = 0.5). If we assume that the true evolutionary history is that all nodes were state 1, Method A has a raw error score of 0.2, whereas method B has a raw error score of 0.125. In other words, we should prefer Method B. If instead, we measure performance using Brier score, Method A scores 0.08, whereas Method B scores 0.125. In other words, we should prefer Method A. This discrepancy is because Brier score applies a quadratic penalty, which is more tolerant of errors with low probability (Supplementary Figure 1A). Consequently, comparing methods with different amounts of uncertainty, or methods in which uncertainty is distributed differently amongst internal nodes, can yield different and even conflicting results depending on whether raw error or Brier score was used to assess performance. This explains discrepancies in the simulation results. For example, applied to symmetrical trees, parsimony is preferred when assessed using raw error (Supplementary Fig. 2A). However, likelihood methods outperform parsimony when assessed using Brier score (Supplementary Fig. 3A).

### Complex likelihood models can be overparameterized

The results of the simulation experiments show that ancestral state estimation using the ARD model is typically less accurate than using the ER model, even when applied to characters simulated using unequal rates. The only experiments in which the ARD model performs better than the ER model occur when characters are simulated under highly unequal transition rates using medium to large trees (≈64-256 tips, Fig 1, Supplementary Figs. 2, 3). The poor performance of the ARD model compared with the ER model is likely a result of the former’s tendency to wildly overestimate transition rates, resulting in higher transition rate estimation error (Supplementary Fig. 6). The ARD model frequently estimates rates many times greater than the true rates under which the data were simulated. In contrast, transition rates estimated under the ER model are typically much closer to the true rates, even when the true rates are unequal (Supplementary Fig. 6). Applying a model with inaccurate transition rates can result in spurious ancestral state estimates. Furthermore, overestimating transition rates will result in more uncertain ancestral state estimates, as states are likely to have changed multiple times on a branch given such a model. This likely explains observed differences in uncertainty between ER and ARD models (Fig. 2, Supplementary Figs. 4, 5).

Inaccurate estimation of transition rates is likely due to overparameterization. The results of the simulation experiments strongly suggest that a single character rarely provides enough information to estimate independent transition rates accurately, particularly when the number of tip states is small. In comparison, assuming transition rates are equal, even when they are not, allows more information to be utilised when estimating transition rates, thus lowering error (Supplementary Fig. 6). This limitation of the ARD model has been highlighted previously (Mooers, A.Ø. and Schluter, D. 1999), although never quantified experimentally. The current study has focussed on binary characters, yet the results likely extend to multistate characters. This is because the number of parameters of the ARD model increases exponentially with the number of states, following *k* = (*n* − 1)^2^ + *n* − 1, where *k* is the number of parameters and n is the number of states. Consequently, for trees of a constant size, increasing the number of states will dramatically decrease the available information from which transition rates can be estimated, resulting in greater transition rate estimation error. Future studies should also consider the symmetrical (SYM) model, which is a less parameter-rich extension of the ER model, in which rates are permitted to vary amongst states, yet forward and backward transition rates are equal (i.e. the model is time reversible) (Harmon, L. 2018). The number of parameters increase with the number of states following *k* = ((*n* − 1)^2^ + *n* − 1)/2. The SYM model is not considered here as it is equivalent to the ER model for binary characters.

Interestingly, applying the true model (i.e. the model under which the data were generated) provides reasonably accurate and certain ancestral state estimates for all experiments irrespective of transition rate heterogeneity (Figs. 2-3, Supplementary Figs. 2-5). This suggests that more accurate estimation of transition rates would reduce ancestral state estimation error and uncertainty under the ARD model. Increasing the sample size (i.e. increasing the number of tips with observed character states) provides additional information for estimating transition rates, thereby increasing accuracy (Salisbury, B.A. and Kim, J. 2001). However, the results presented here suggest that above ≈128 tips, increasing the sample size provides diminishing returns (Supplementary Fig. 6). An alternative solution would be to better constrain rate estimates. This can be achieved using a traditional Bayesian approach, whereby the transition rate prior is specified, rather than estimated from the data. Such an approach is already implemented in the popular phylogenetic software package revBayes (Höhna, S., Landis, M.J., et al. 2016). One potential drawback of this approach is that, without any prior knowledge of reasonable transition rates, the specified prior would have to be diffuse. As such, it may not preclude transition rate overestimation. Alternatively, the transition rate prior could be estimated from multiple characters, rather than a single character. A drawback of this approach is that states are not equivalent in morphological or ecological characters, unlike molecular characters. Character transitions do not necessarily share a common evolutionary process. However, empirical morphological datasets have been demonstrated to exhibit similar exponential distributions of homoplasy, with many low homoplasy characters and few high homoplasy characters (Goloboff, P.A., Torres, A., et al. 2017, Murphy, J.L., Puttick, M.N., et al. 2021). It therefore seems reasonable to assume that transition rates of morphological characters, and perhaps even ecological characters, share a similar distribution. Regardless of whether the transition rate prior is estimated from multiple characters or specified a priori, constraining transition rates to avoid overestimation should improve performance of the ARD model.

### Difference in uncertainty is a better predictor of model performance than difference in likelihood

One of the most surprising results of the current study is that difference in model performance is better predicted by difference in uncertainty between ancestral state estimates than by difference in model likelihood. As well as having a greater diagnostic power (see Table 1), difference in uncertainty between two models is proportional to difference in error (Fig. 6), which is a prerequisite for model averaging. In contrast, difference in AIC or BIC appears to be inversely proportional to difference in error, which is the exact opposite of the expected behaviour (Fig. 7, Supplementary Fig. 21). This suggests that model averaging using likelihood-based information criteria such as AIC or BIC will result in giving maximum weight to the least accurate model. One possible explanation of this strange result is that models with unrealistically high transition rates are frequently assigned high likelihood. Preventing overestimation of transition rates (see above) may go some way to correcting this problem. Until this is addressed, however, it is suggested that ancestral state estimation models should be selected and averaged using uncertainty, while likelihood-based information criteria should be avoided.

### Understanding how tree properties affect ancestral state estimation informs sampling strategy

The results presented here highlight how data properties such as tree size, shape, ultrametricity, branch length heterogeneity, transition rate and rate heterogeneity affect ancestral state estimation. Importantly some of these properties, namely those related to the phylogeny, are a product of both the evolutionary history of the data and sampling strategy imposed by the researcher. Consequently, sampling strategy to some extent determines tree properties and thus the accuracy of ancestral state estimates. The results of the simulation study demonstrate that increasing the sample size (i.e. the number of observed character states) reduces the error of ancestral state estimates, regardless of the method used. However, the relationship is nonlinear; increasing the sample size provides diminishing returns as tree size increases (Fig 1, Supplementary Figs. 2, 3). The simulation results are equivocal about the effect of tree shape. In principle, asymmetrical trees should provide more accuracy than symmetrical trees, as they have fewer nested internal nodes (i.e. internal nodes which parent only internal nodes). As such, tip states more directly inform ancestral state estimates of internal nodes. However, this ignores the effect of branch lengths. Trees with long branches will produce less accurate ancestral estimates, and this can override any benefit of tree asymmetry (Mooers, A.Ø. 2004). The results presented in the current study suggest that ancestral state estimates are more accurate when applied to trees lacking long branches or substantial branch length heterogeneity. Long branches can be mitigated by inclusion of fossils. Fossils break long branches (Cobbett, A., Wilkinson, M., et al. 2007), resulting in non-ultrametric trees. They are also closer to internal nodes of the tree than extant taxa. As such, trees that include fossils tend to exhibit reduced average branch lengths and branch length heterogeneity compared to corresponding trees without fossils. Previous studies have already demonstrated that inclusion vs. exclusion of fossils can yield substantively different ancestral estimates (Finarelli, J.A. and Flynn, J.J. 2006, Puttick, M.N. 2016). The simulation results presented here suggest that including fossils in extant-only trees is likely to improve accuracy, irrespective of the method used to estimate ancestral states. Conversely, employing a sampling strategy that excludes fossils will likely limit accuracy, irrespective of how many taxa are sampled.

## Conclusion

The Equal Rates (ER) model is the most accurate over most simulations. It performs well over a range of tree shapes and generally outperforms the All Rates Different (ARD) model, even when the data are generated under highly unequal rates. Different methods perform better or worse depending on the properties of the data, as well as the metric used to quantify error. Parsimony performs poorly when applied to trees with long branches. The ARD model frequently overestimates transition rates indicating the model is overparameterized; there is not enough information in the data to accurately estimate independent transition rates between states. Comparing model selection approaches reveals a surprising result: Difference in log-likelihood based information criteria is a poor predictor of difference in model error, whereas difference in the uncertainty of ancestral state estimates predicts difference in model error reasonably well. Thus, models with more certain ancestral state estimates tend to be more accurate. Based on the results presented here, I propose the following recommendations for future ancestral state estimation studies:

1. Sample as many tips as possible This reduces error of ancestral state and transition rate estimates. If possible, sample more than ≈128 tips, as sample sizes smaller than this are much more error prone.
2. Include fossils if possible. Fossils break long branches, thereby lowering the average branch length and branch length heterogeneity of extant-only trees. This, in turn, will reduce the error of ancestral state and transition rate estimates.
3. Select models based on the difference in uncertainty of ancestral estimates rather than difference in log-likelihood scores. Log-likelihood based metrics such as ΔAIC and ΔBIC have less diagnostic power than Δuncertainty for predicting the least erroneous ancestral state estimation model.
4. Benchmark ancestral state estimates using the equal rates (ER) model. ER estimates of ancestral states and transition rates are fairly accurate and seem to be robust to model violation. Consequently, if estimates using alternative methods differ from those of the ER model, their adequacy should be critically considered.

## Acknowledgements

I thank Phil Donoghue, Mike Benton, Russell Garwood, Pierre Cockx (University of Bristol) and Martin Smith (Durham University) for their extremely useful feedback. Computational analyses were conducted using the Brown Nugget high-performance workstation at the University of Bristol. The research was made possible through support of the ERC grant no. 788203 (INNOVATION).

