## Supplementary Figures for "What is the best method for estimating ancestral states from discrete characters?" (from another version of this preprint: version 1, 2023-09-01)

**J. N. Keating**

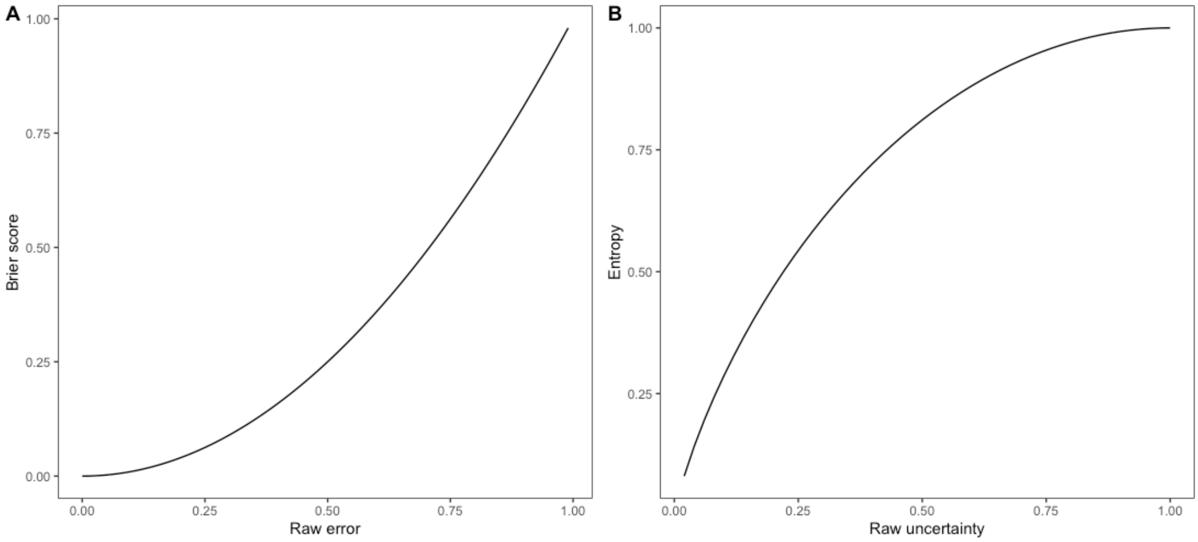


***S. Figure 1.*** *Comparison of raw and Brier measures of estimation error (**A**); Comparison of raw and Shannon entropy measures of estimation uncertainty (**B**).*

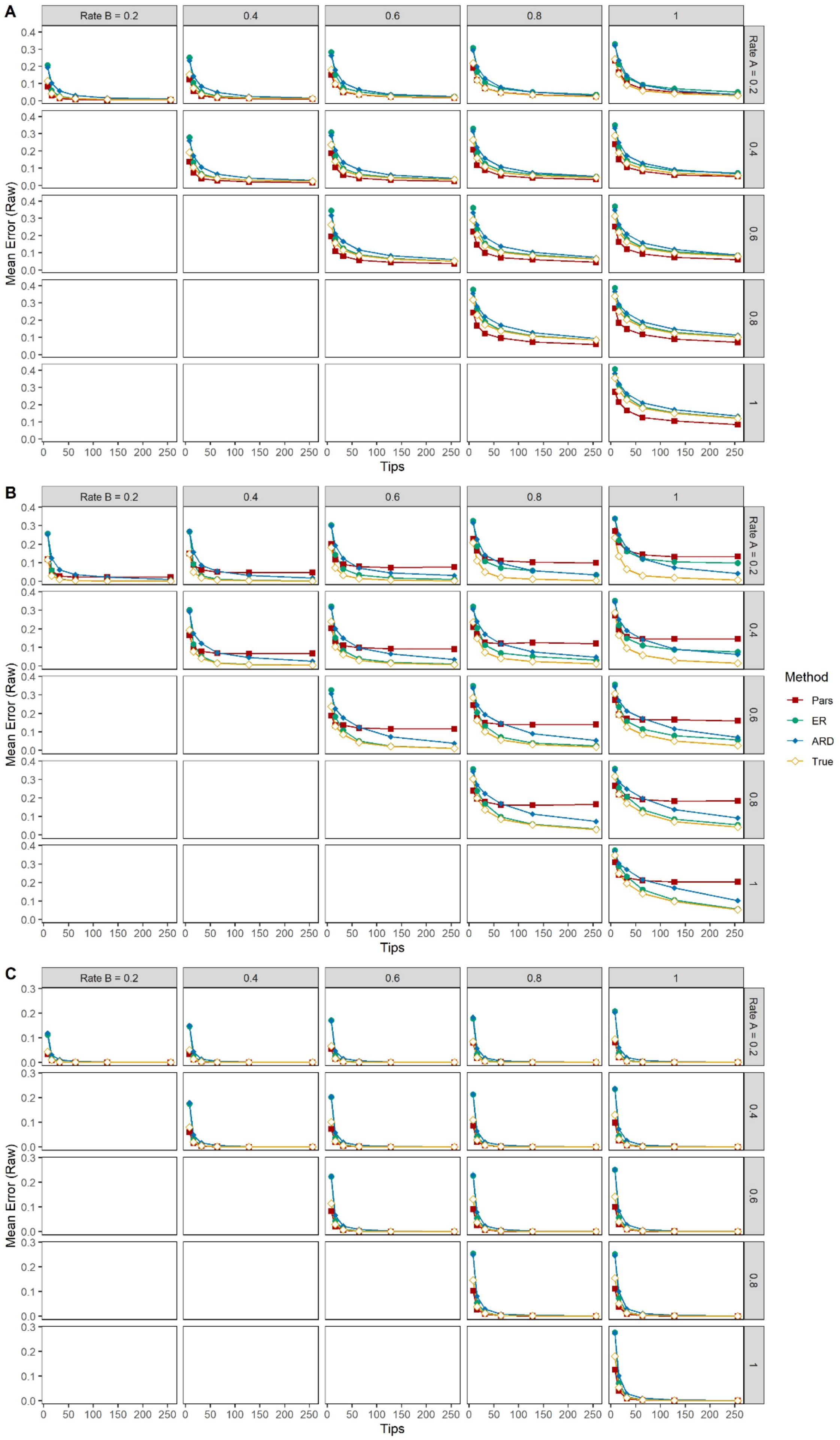
A
B
C
Rate B = 0.2
0.4
0.6
0.8
1
Rate A = 0.2
Mean Error (Raw)
Tips
Method
Pars
ER
ARD
True

***S. Figure 2.*** *Mean raw error of different ancestral states estimation methods plotted against the number of tips present in the tree used to simulate character data: ACCTRAN Parsimony (Pars); Equal Rates model (ER); All-Rates-Different model (ARD); the true model under which the data were simulated (True). Results are presented for 15 different generating models and three different tree shapes: symmetrical equal branch length (**A**); asymmetrical ultrametric (**B**); and asymmetrical equal branch length (**C**). Each data point represents the mean of 500 simulated characters. Error bars show the 95% confidence interval of the mean. All methods exhibit less error when more tips are present. ARD typically has higher mean error than all other methods. Parsimony has very low mean error when branch lengths are equal, but is outperformed by likelihood models when branch lengths are unequal.*

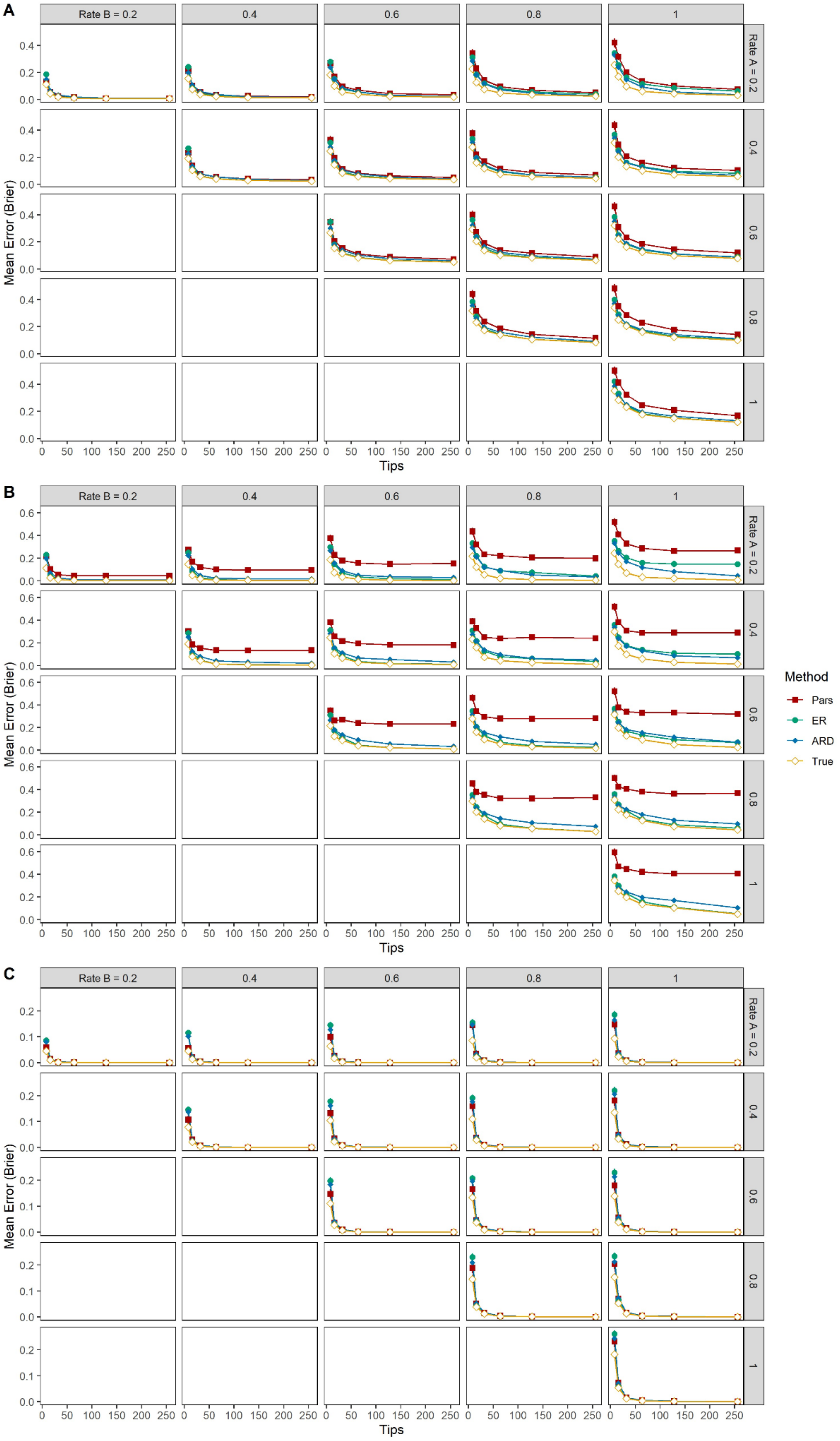
A
B
C
Rate B = 0.2
0.4
0.6
0.8
1
Rate A = 0.2
0.4
0.6
0.8
1
Mean Error (Brier)
Tips
0 50 100 150 200 250
Method
Pars
ER
ARD
True

***S. Figure 3.*** *Mean Brier error of different ancestral states estimation methods plotted against the number of tips present in the tree used to simulate character data: ACCTRAN Parsimony (Pars); Equal Rates model (ER); All-Rates-Different model (ARD); the true model under which the data were simulated (True). Results are presented for 15 different generating models and three different tree shapes: symmetrical equal branch length (**A**); asymmetrical ultrametric (**B**); and asymmetrical equal branch length (**C**). Each data point represents the mean of 500 simulated characters. Error bars show the 95% confidence interval of the mean. All methods exhibit less error when more tips are present. ARD typically has higher mean error than all other methods. Parsimony has very low mean error when branch lengths are equal, but is outperformed by likelihood models when branch lengths are unequal.*

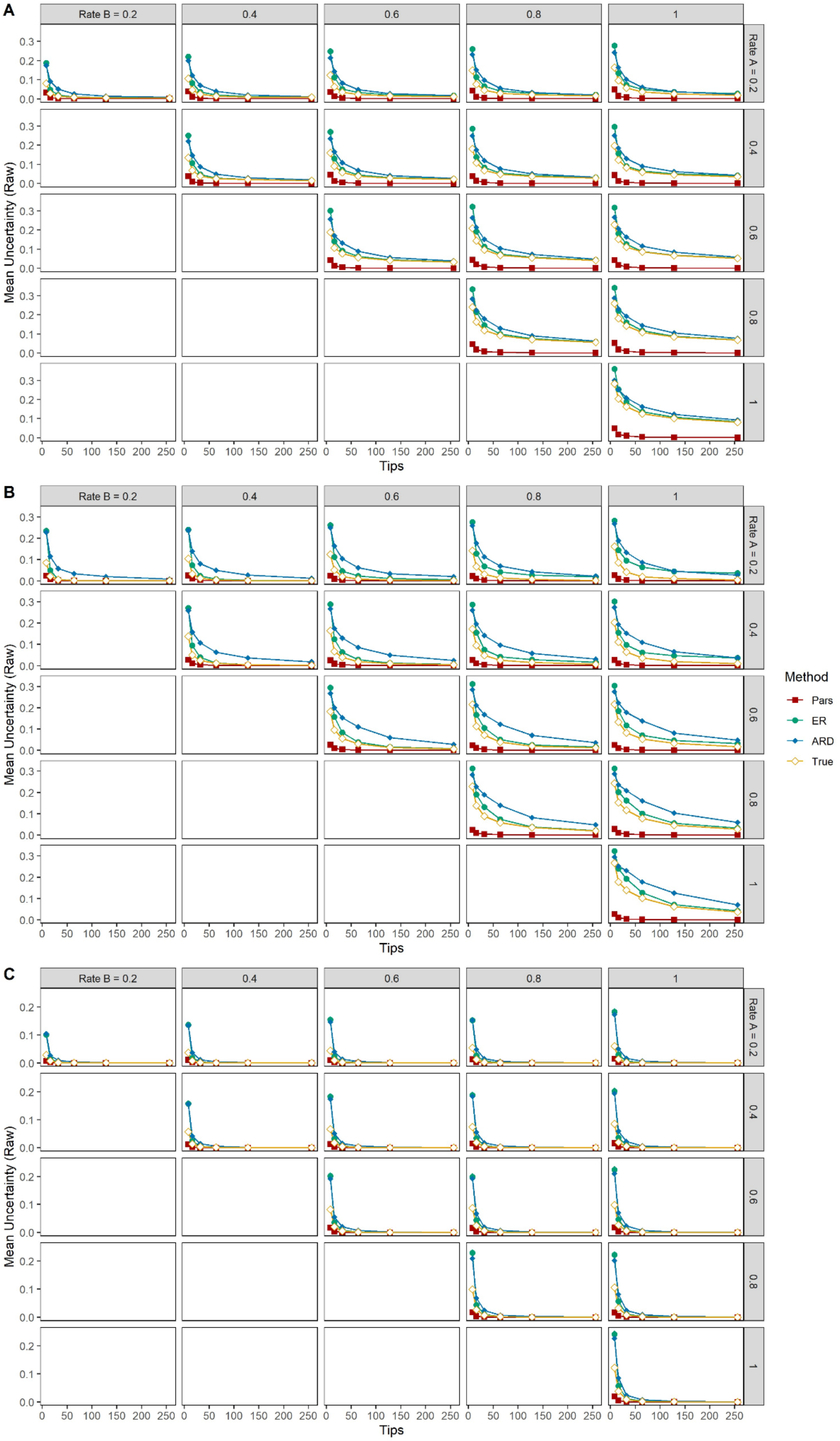
A
B
C
Rate B = 0.2
0.4
0.6
0.8
1
Rate A = 0.2
0.4
0.6
0.8
1
Mean Uncertainty (Raw)
Tips
0 50 100 150 200 250
0.0 0.1 0.2 0.3
Method
Pars
ER
ARD
True

***S. Figure 4.*** *Mean raw uncertainty of different ancestral states estimation methods plotted against the number of tips present in the tree used to simulate character data: ACCTRAN Parsimony (Pars); Equal Rates model (ER); All-Rates-Different model (ARD); the true model under which the data were simulated (True). Results are presented for 15 different generating models and three different tree shapes: symmetrical equal branch length (**A**); asymmetrical ultrametric (**B**); and asymmetrical equal branch length (**C**). Each data point represents the mean of 500 simulated characters. Error bars show the 95% confidence interval of the mean. All methods exhibit less uncertainty when more tips are present. ARD typically has higher mean uncertainty, while parsimony exhibits lower mean uncertainty than all other methods.*

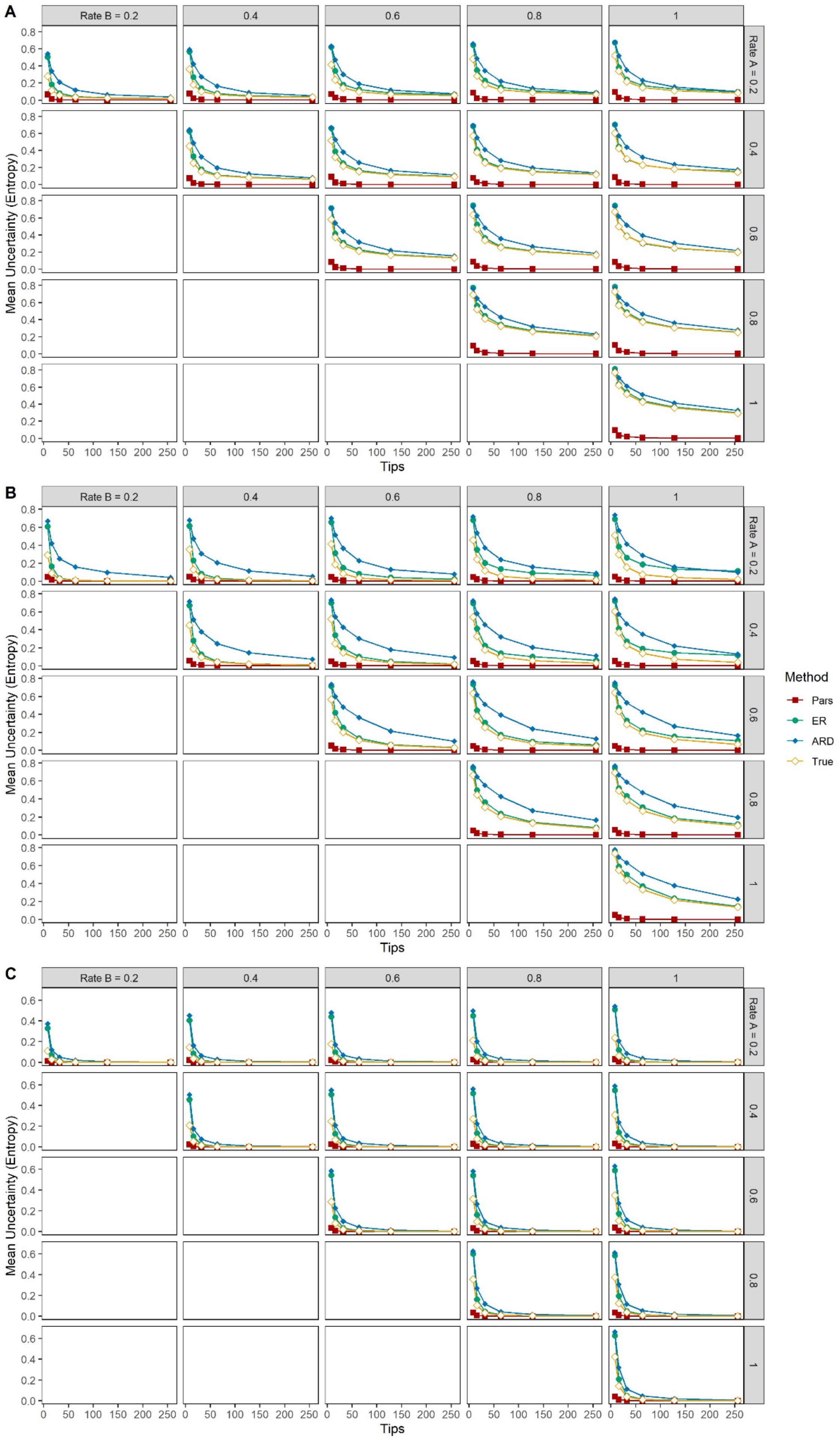
A
B
C
Rate B = 0.2
0.4
0.6
0.8
1
Rate A = 0.2
0.4
0.6
0.8
1
Mean Uncertainty (Entropy)
Tips
0
50
100
150
200
250
0.0
0.2
0.4
0.6
0.8
Method
Pars
ER
ARD
True

***S. Figure 5.*** *Mean entropy of different ancestral states estimation methods plotted against the number of tips present in the tree used to simulate character data: ACCTRAN Parsimony (Pars); Equal Rates model (ER); All-Rates-Different model (ARD); the true model under which the data were simulated (True). Results are presented for 15 different generating models and three different tree shapes: symmetrical equal branch length (**A**); asymmetrical ultrametric (**B**); and asymmetrical equal branch length (**C**). Each data point represents the mean of 500 simulated characters. Error bars show the 95% confidence interval of the mean. All methods exhibit less uncertainty when more tips are present. ARD typically has higher mean uncertainty, while parsimony exhibits lower mean uncertainty than all other methods.*

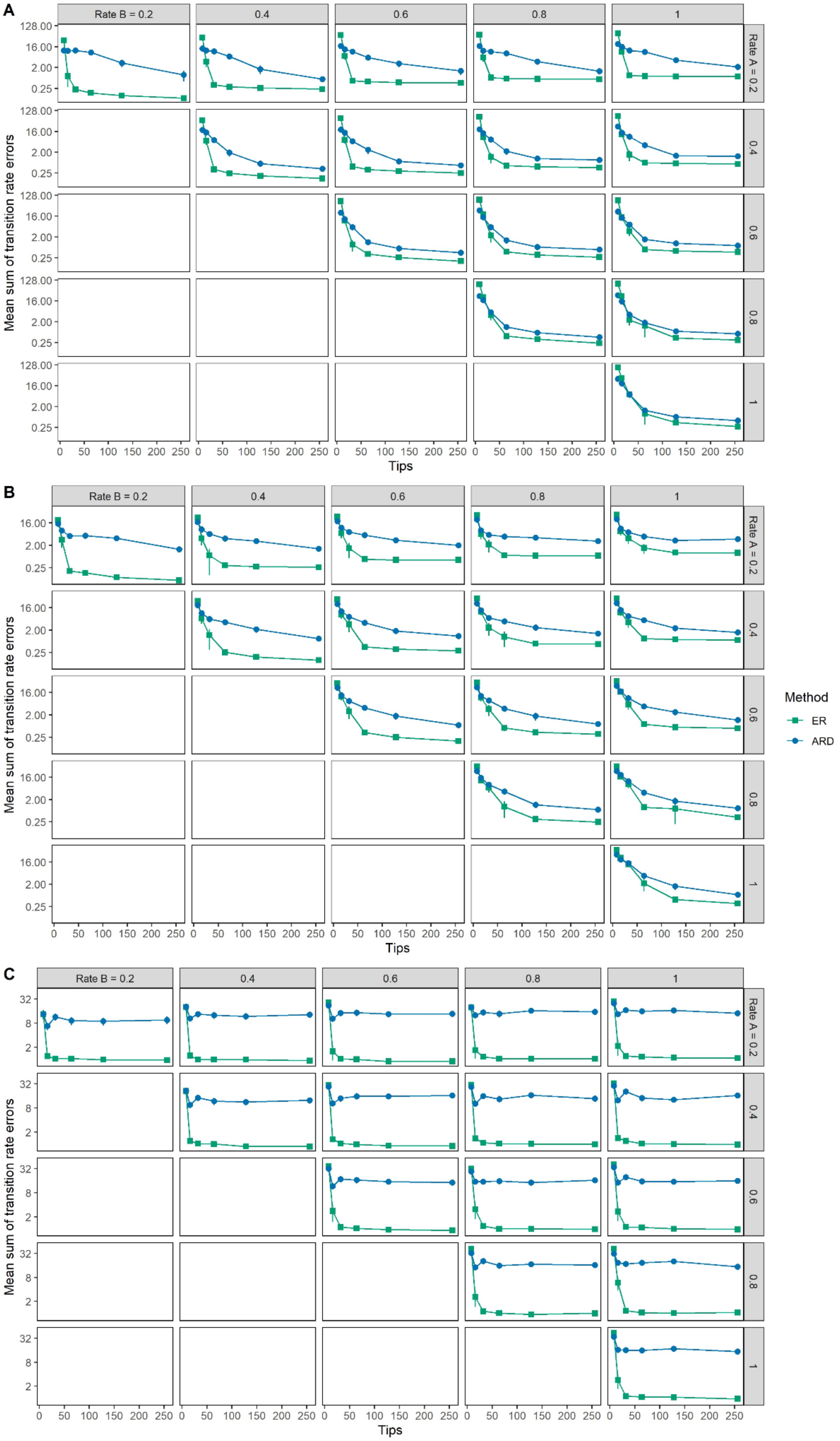
A
B
C
Rate B = 0.2
0.4
0.6
0.8
1
Rate A = 0.2
0.4
0.6
0.8
1
128.00
16.00
2.00
0.25
32
8
2
0
50
100
150
200
250
Tips
Mean sum of transition rate errors
Method
ER
ARD

***S. Figure 6.*** *Mean sum of estimated transition rates of Equal Rates (ER) and All Rates Different (ARD) models plotted against the number of tips present in the tree used to simulate character data. Results are presented for 15 different generating models and three different tree shapes: symmetrical equal branch length (**A**); asymmetrical ultrametric (**B**); and asymmetrical equal branch length (**C**). Each data point represents the mean of 500 simulated characters. Error bars show the 95% confidence interval of the mean. For most experiments the ARD model tends to overestimate transition rates. Increasing the number of tips tends to mitigate overestimation of transition rates for both ER and ARD models. An exception is observed when the ARD model is applied to an asymmetrical tree with equal branch lengths. Here, the sum of estimated transition rates increases with the number of tips.*

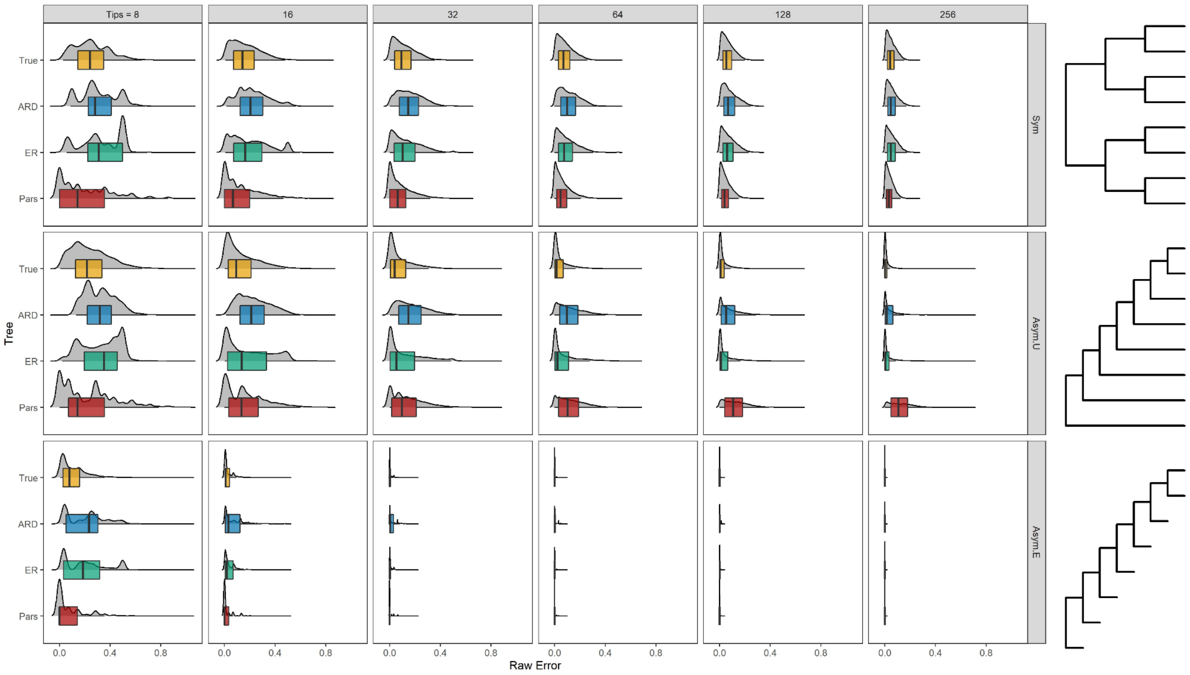


***S. Figure 7.*** *Overall distributions of raw error, per tree topology. Each boxplot summarises the results of 7500 binary characters generated under 15 different transition models using one of three tree topologies: symmetrical equal branch lengths (Sym); asymmetrical ultrametric (Asym.U); asymmetrical equal branch lengths (Asym.E). Ancestral states were estimated using ACCTRAN Parsimony (Pars); Equal Rates model (ER); All-Rates-Different model (ARD); the true model under which the data were generated (True). All methods exhibit less error when the tree is asymmetrical with equal branch lengths. Likelihood models tend to show marginally higher uncertainty when applied to symmetrical vs. asymmetrical trees. Parsimony shows the opposite pattern; higher uncertainty when applied to asymmetrical vs. symmetrical trees.*

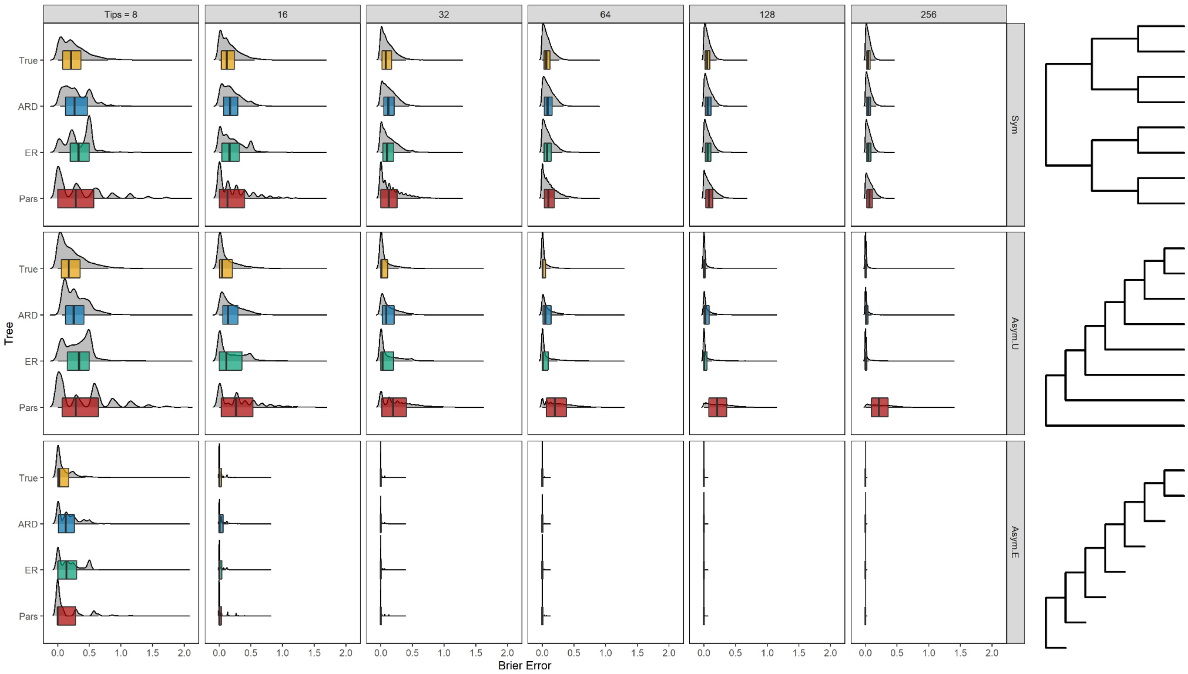


***S. Figure 8.*** *Overall distributions of Brier error, per tree topology. Each boxplot summarises the results of 7500 binary characters generated under 15 different transition models using one of three tree topologies: symmetrical equal branch lengths (Sym); asymmetrical ultrametric (Asym.U); asymmetrical equal branch lengths (Asym.E). Ancestral states were estimated using ACCTRAN Parsimony (Pars); Equal Rates model (ER); All-Rates-Different model (ARD); the true model under which the data were generated (True). All methods exhibit less error when the tree is asymmetrical with equal branch lengths. Likelihood models show similar levels of uncertainty when applied to asymmetrical vs. symmetrical trees. Parsimony shows higher uncertainty when applied to asymmetrical vs. symmetrical trees.*

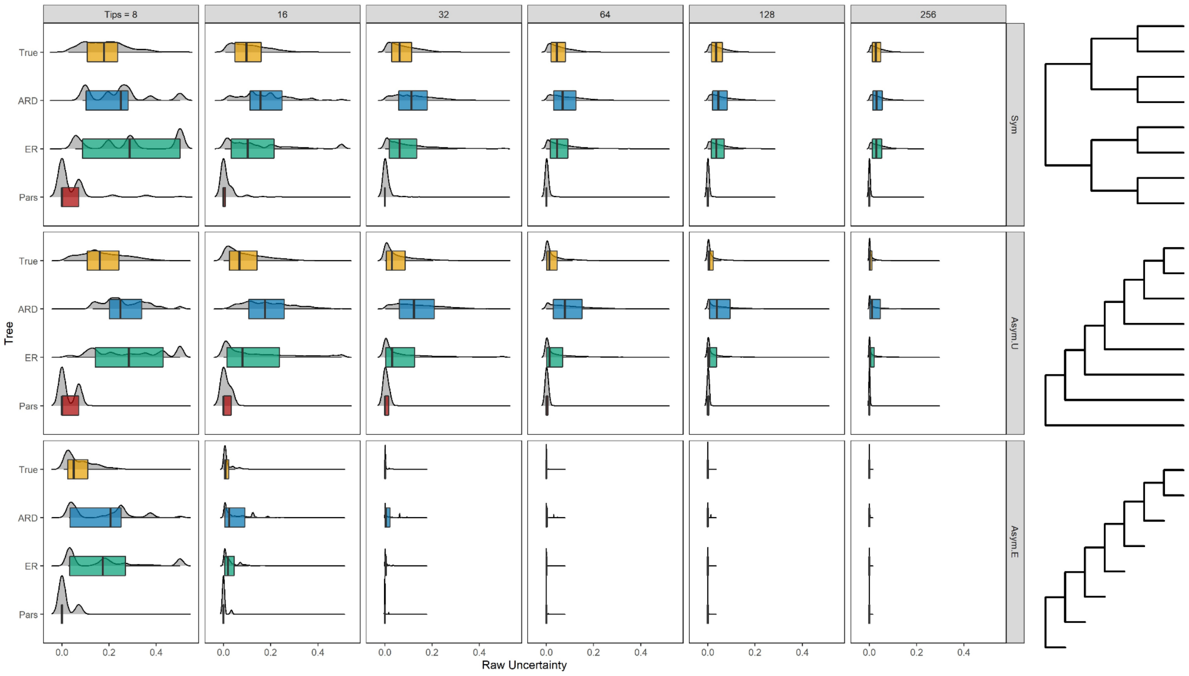


***S. Figure 9.*** *Overall distributions of raw uncertainty, per tree topology. Each boxplot summarises the results of 7500 binary characters generated under 15 different transition models using one of three tree topologies: symmetrical equal branch lengths (Sym); asymmetrical ultrametric (Asym.U); asymmetrical equal branch lengths (Asym.E). Ancestral states were estimated using ACCTRAN Parsimony (Pars); Equal Rates model (ER); All-Rates-Different model (ARD); the true model under which the data were generated (True). Parsimony consistently exhibits the least uncertainty. All methods exhibit less uncertainty when the tree is asymmetrical with equal branch lengths. Likelihood models exhibit marginally less uncertainty when applied to asymmetrical vs. symmetrical ultrametric trees.*

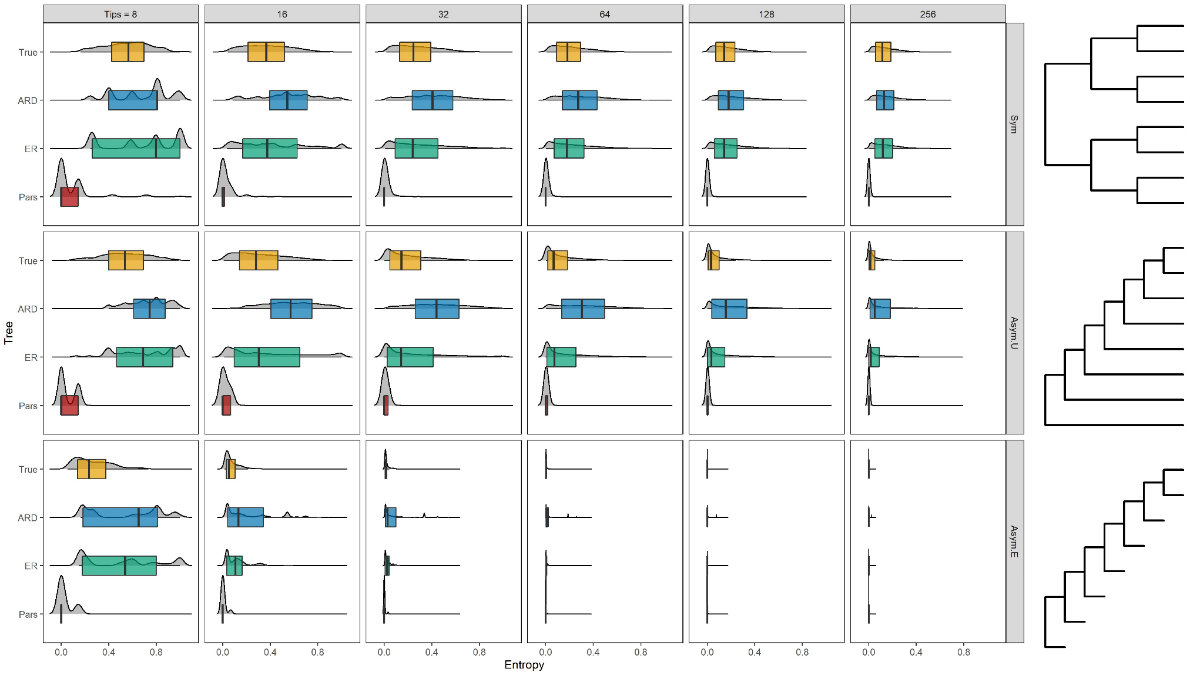


***S. Figure 10.*** *Overall distributions of entropy, per tree topology. Each boxplot summarises the results of 7500 binary characters generated under 15 different transition models using one of three tree topologies: symmetrical equal branch lengths (Sym); asymmetrical ultrametric (Asym.U); asymmetrical equal branch lengths (Asym.E). Ancestral states were estimated using ACCTRAN Parsimony (Pars); Equal Rates model (ER); All-Rates-Different model (ARD); the true model under which the data were generated (True). Parsimony consistently exhibits the least uncertainty. All methods exhibit less entropy when the tree is asymmetrical with equal branch lengths. Likelihood methods exhibit marginally less entropy when applied to asymmetrical vs. symmetrical ultrametric trees.*

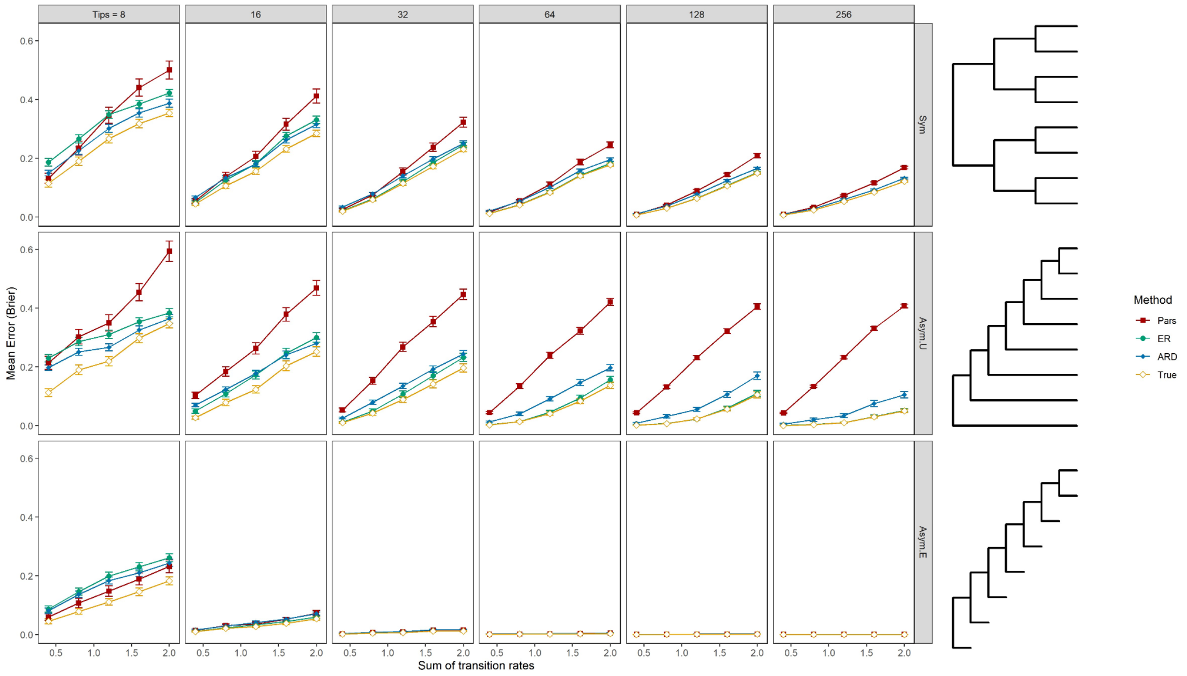

***S. Figure 11.*** *Mean Brier error of different ancestral states estimation methods plotted against the sum of transition rates used to simulate character data: ACCTRAN Parsimony (Pars); Equal Rates model (ER); All-Rates-Different model (ARD); the true model under which the data were simulated (True). Each data point represents the mean of 500 binary characters generated under an ER model. Error bars show the 95% confidence interval of the mean. Results are plotted for trees of size 8-256 tips using three topologies: symmetrical equal branch lengths (Sym); asymmetrical ultrametric (Asym.U); asymmetrical equal branch lengths (Asym.E). All methods exhibit less error when the sum of transition rates is lower.*

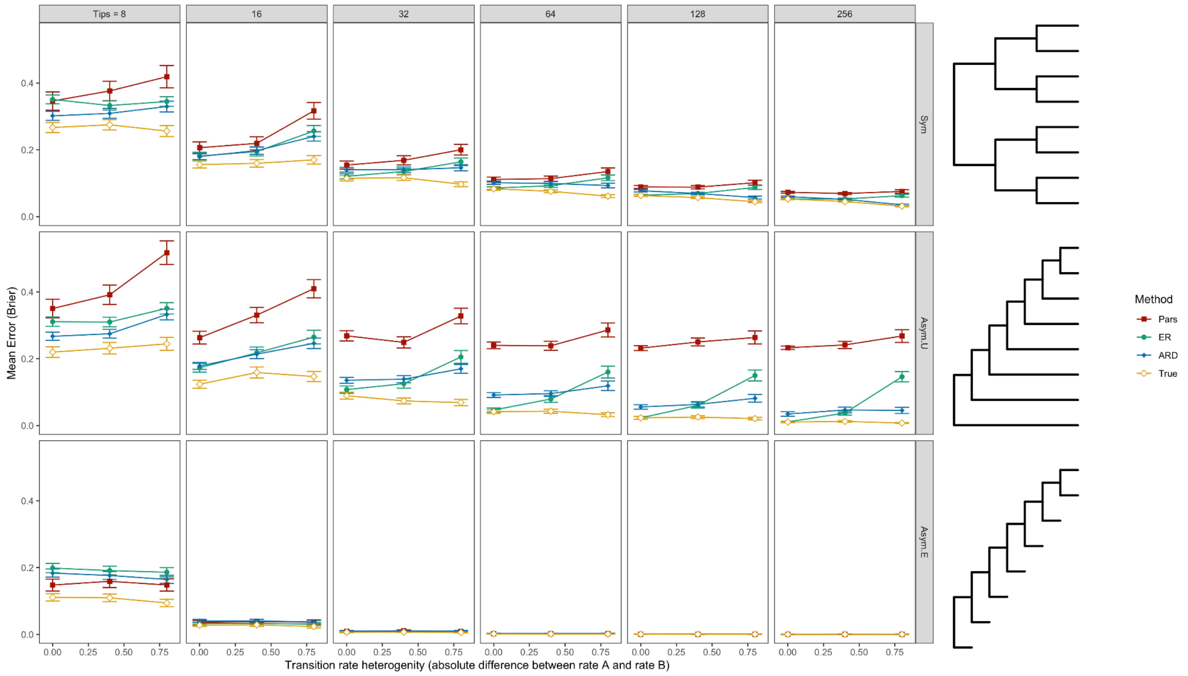


***S. Figure 12.*** *Mean Brier error of different ancestral states estimation methods plotted against the transition rate heterogeneity used to simulate character data, calculated by taking the absolute difference between forward and backward transition rates. ACCTRAN Parsimony (Pars); Equal Rates model (ER); All-Rates-Different model (ARD); the true model under which the data were simulated (True). Each data point represents the mean of 500 binary characters generated under a model in which the sum of transition rates is equal to 1.2. Error bars show the 95% confidence interval of the mean. Results are plotted for trees of size 8-256 tips using three topologies: symmetrical equal branch lengths (Sym); asymmetrical ultrametric (Asym.U); asymmetrical equal branch lengths (Asym.E). Using ultrametric trees, ER and parsimony error tends to marginally increase as transition rate heterogeneity increases.*

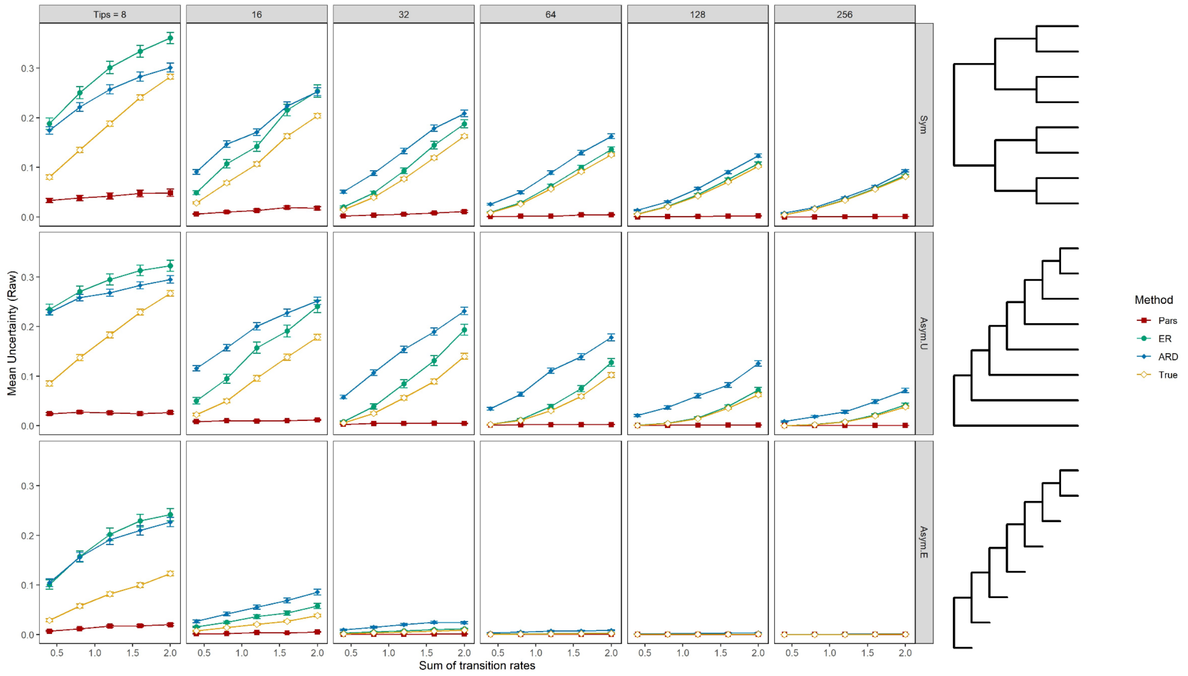


***S. Figure 13.*** *Mean raw uncertainty of different ancestral states estimation methods plotted against the sum of transition rates used to simulate character data: ACCTRAN Parsimony (Pars); Equal Rates model (ER); All-Rates-Different model (ARD); the true model under which the data were simulated (True). Each data point represents the mean of 500 binary characters generated under a model in which the sum of transition rates is equal to 1.2. Error bars show the 95% confidence interval of the mean. Results are plotted for trees of size 8-256 tips using three topologies: symmetrical equal branch lengths (Sym); asymmetrical ultrametric (Asym.U); asymmetrical equal branch lengths (Asym.E). Parsimony consistently shows very low uncertainty. All likelihood models show increasing uncertainty as the sum of transition rates increases. An exception can be seen in large asymmetrical trees with equal branch lengths, where raw uncertainty remains low for the ER model, as well as the true model under which the data were generated.*

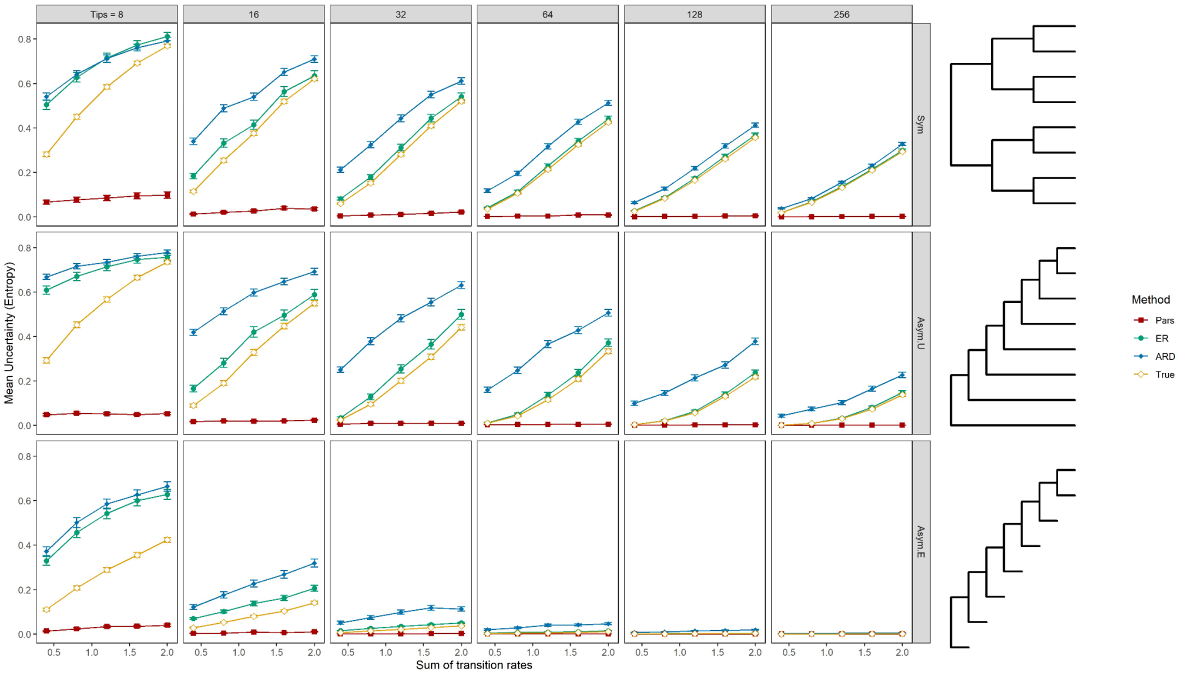


***S. Figure 14.*** *Mean Entropy of different ancestral states estimation methods plotted against the sum of transition rates used to simulate character data: ACCTRAN Parsimony (Pars); Equal Rates model (ER); All-Rates-Different model (ARD); the true model under which the data were simulated (True). Each data point represents the mean of 500 binary characters generated under a model in which the sum of transition rates is equal to 1.2. Error bars show the 95% confidence interval of the mean. Results are plotted for trees of size 8-256 tips using three topologies: symmetrical equal branch lengths (Sym); asymmetrical ultrametric (Asym.U); asymmetrical equal branch lengths (Asym.E). Parsimony consistently shows very low entropy. All likelihood methods show increasing entropy as the sum of transition rates increases. An exception can be seen in large asymmetrical trees with equal branch lengths, where entropy remains low for the ER model, as well as the true model under which the data were generated.*

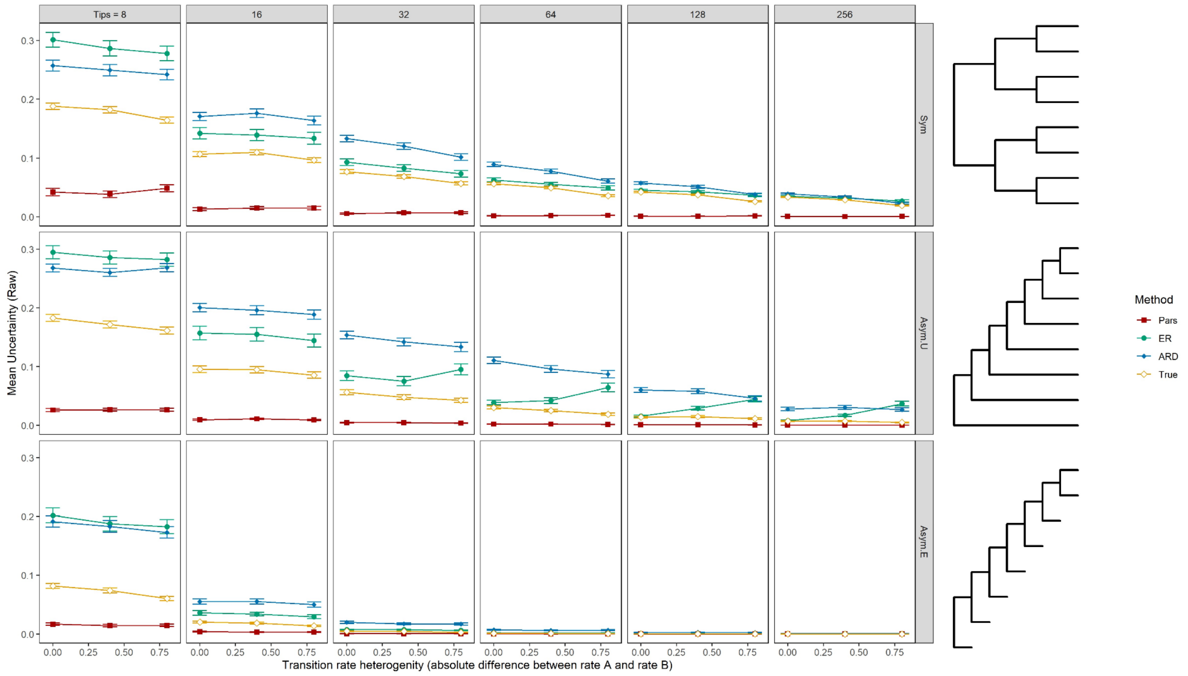


***S. Figure 15.*** *Mean raw uncertainty of different ancestral states estimation methods plotted against the transition rate heterogeneity used to simulate character data: ACCTRAN Parsimony (Pars); Equal Rates model (ER); All-Rates-Different model (ARD); the true model under which the data were simulated (True). Each data point represents the mean of 500 binary characters generated under a model in which the sum of transition rates is equal to 1.2. Error bars show the 95% confidence interval of the mean. Results are plotted for trees of size 8-256 tips using three topologies: symmetrical equal branch lengths (Sym); asymmetrical ultrametric (Asym.U); asymmetrical equal branch lengths (Asym.E). Parsimony consistently shows very low uncertainty. Likelihood models tend to show a slight decrease in uncertainty as rate heterogeneity increases. One notable exception is the ER model applied to asymmetrical trees of 32 tips or greater, which shows uncertainty increasing with transition rate heterogeneity.*

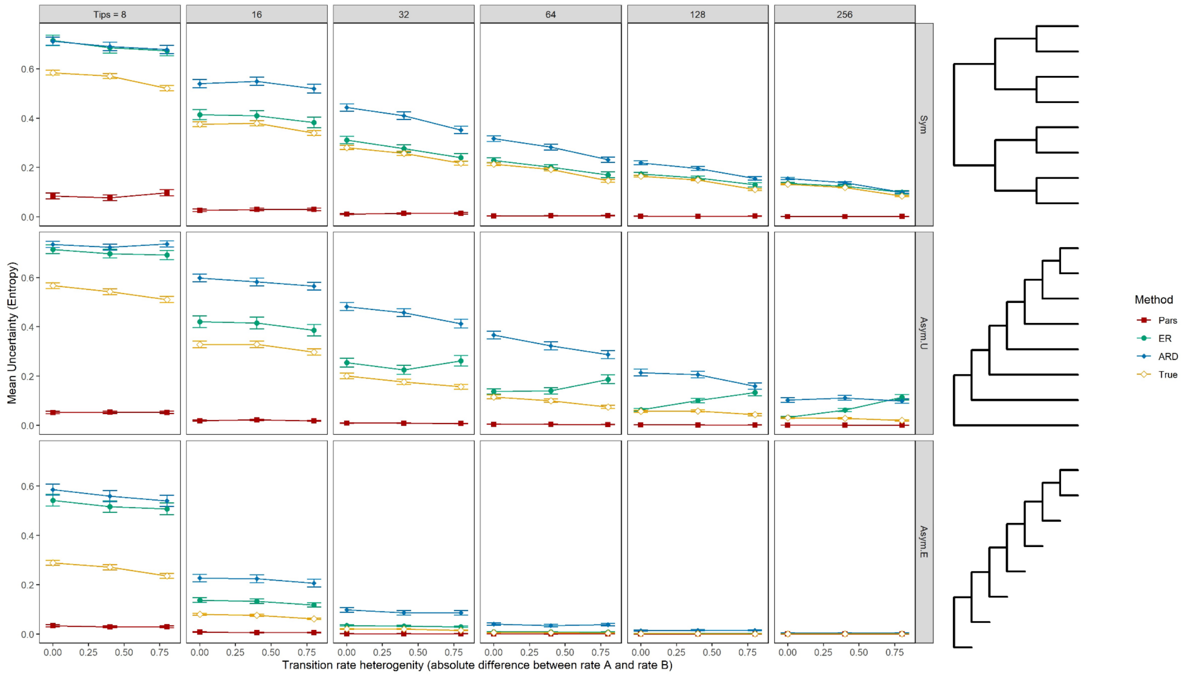


***S. Figure 16.*** *Mean Brier uncertainty of different ancestral states estimation methods plotted against the transition rate heterogeneity used to simulate character data: ACCTRAN Parsimony (Pars); Equal Rates model (ER); All-Rates-Different model (ARD); the true model under which the data were simulated (True). Each data point represents the mean of 500 binary characters generated under a model in which the sum of transition rates is equal to 1.2. Error bars show the 95% confidence interval of the mean. Results are plotted for trees of size 8-256 tips using three topologies: symmetrical equal branch lengths (Sym); asymmetrical ultrametric (Asym.U); asymmetrical equal branch lengths (Asym.E). Parsimony consistently shows very low uncertainty. Likelihood models tend to show a slight decrease in uncertainty as rate heterogeneity increases. One notable exception is the ER model applied to asymmetrical trees of 32 tips or greater, which shows uncertainty increasing with transition rate heterogeneity.*

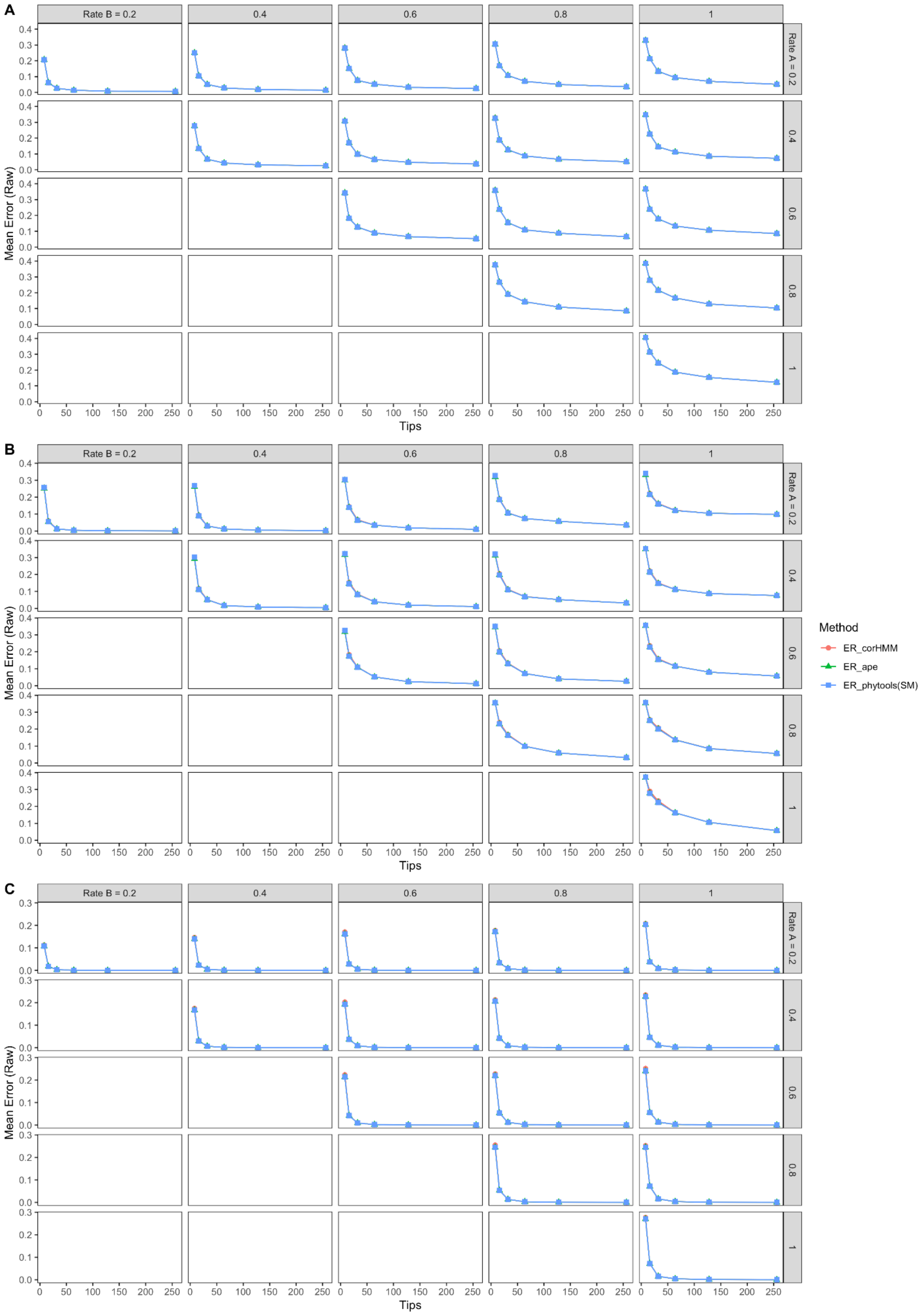


***S. Figure 17.*** *Mean raw error of the ER model implemented using different likelihood optimisation methods plotted against the number of tips present in the tree used to simulate character data: Marginal likelihood estimation using corHMM (ARD_corHMM); scaled*

*conditional likelihood using ape (ARD_ape); stochastic mapping using phytools (ARD_phytools(SM)). Results are presented for 15 different generating models and three different tree shapes: symmetrical equal branch length (**A**); asymmetrical ultrametric (**B**); and asymmetrical equal branch length (**C**). Each data point represents the mean of 500 simulated characters. Error bars show the 95% confidence interval of the mean. All likelihood optimisation methods using the ER model exhibit identical raw error.*

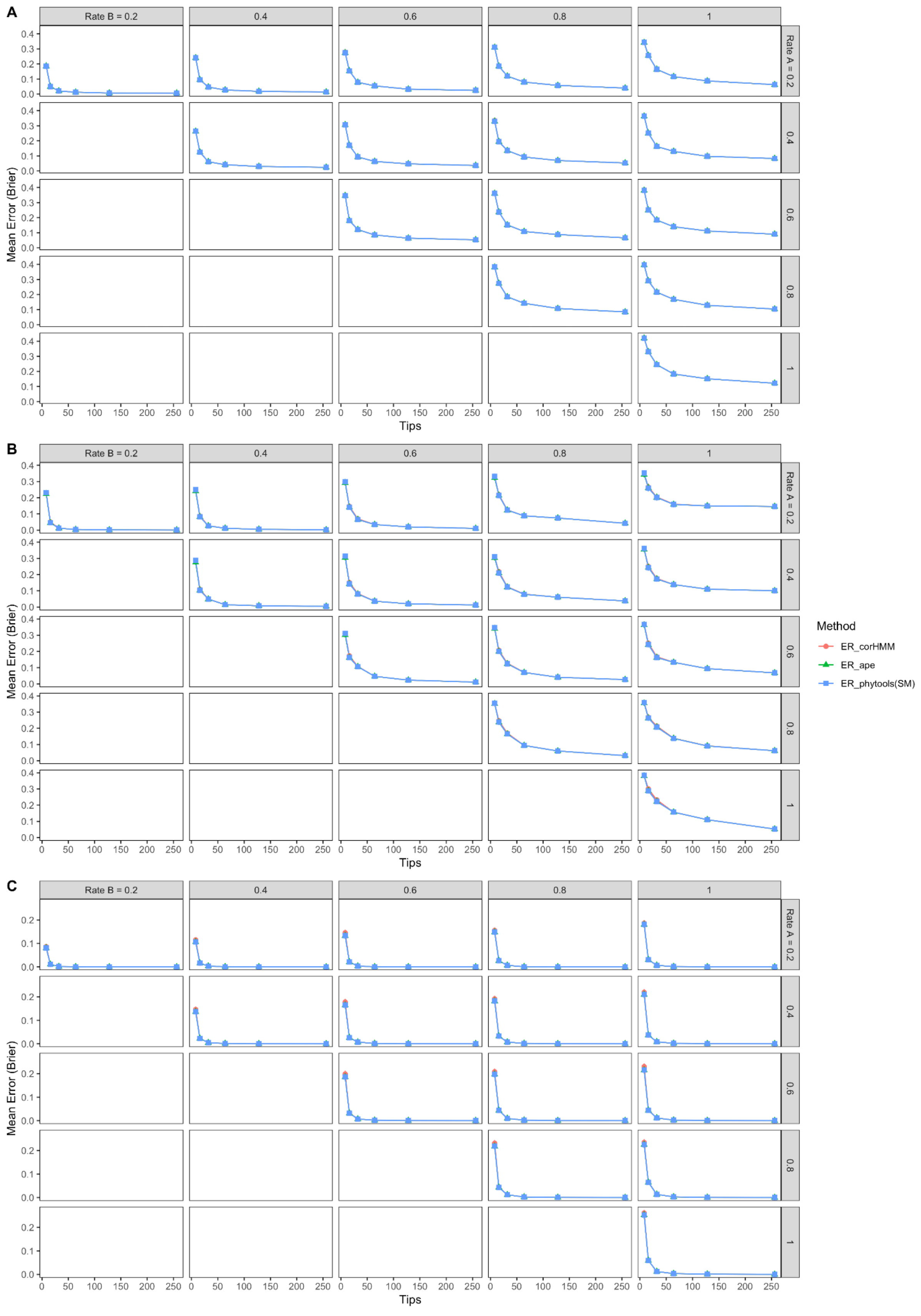


***S. Figure 18.*** *Mean Brier error of the ER model implemented using different likelihood optimisation methods plotted against the number of tips present in the tree used to simulate character data: Marginal likelihood estimation using corHMM (ARD_corHMM); scaled*

*conditional likelihood using ape (ARD_ape); stochastic mapping using phytools (ARD_phytools(SM)). Results are presented for 15 different generating models and three different tree shapes: symmetrical equal branch length (**A**); asymmetrical ultrametric (**B**); and asymmetrical equal branch length (**C**). Each data point represents the mean of 500 simulated characters. Error bars show the 95% confidence interval of the mean. All likelihood optimisation methods using the ER model exhibit identical Brier error.*

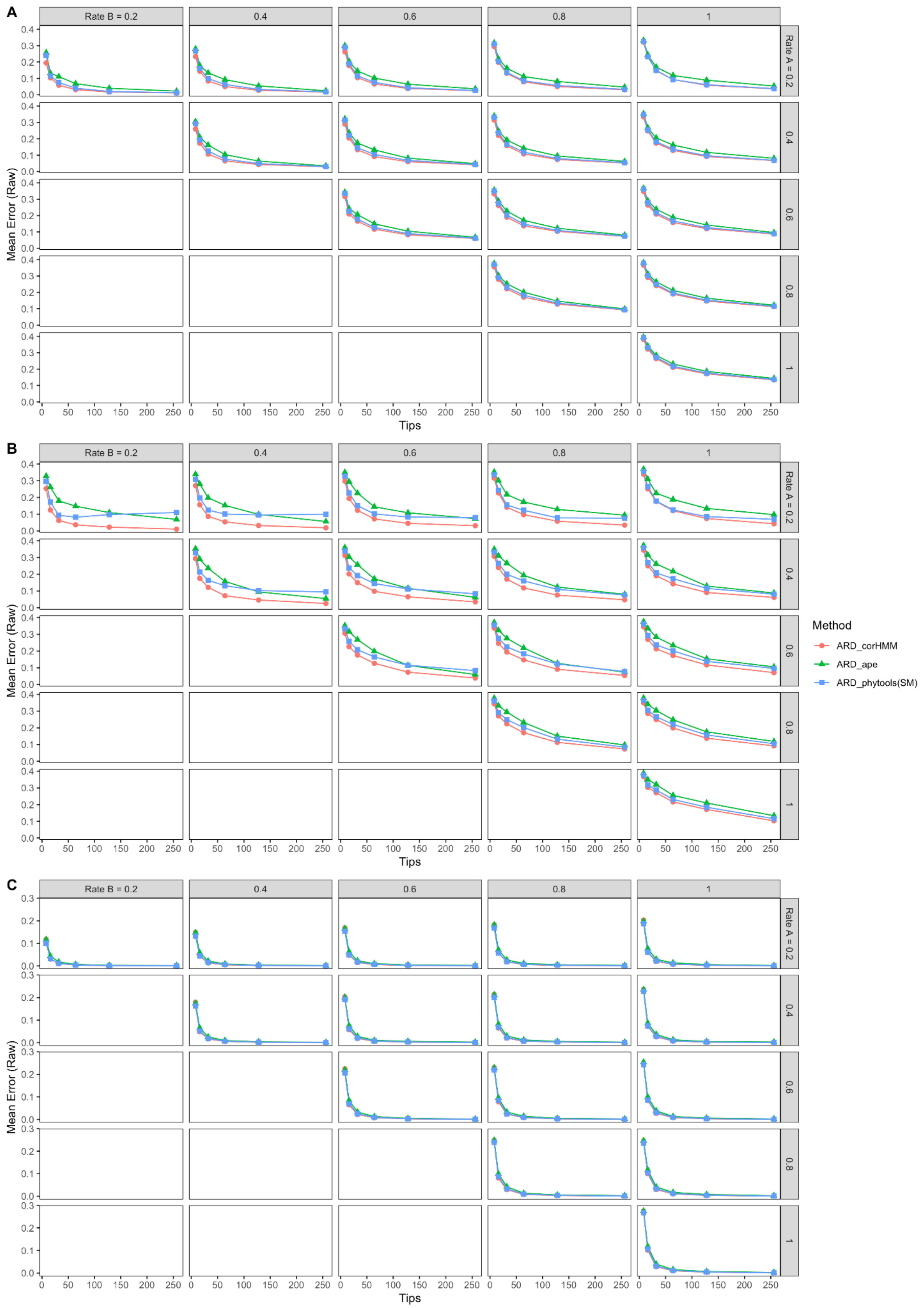


***S. Figure 19.*** *Mean raw error of the ARD model implemented using different likelihood optimisation methods plotted against the number of tips present in the tree used to simulate character data: Marginal likelihood estimation using corHMM (ARD_corHMM); scaled*

*conditional likelihood using ape (ARD_ape); stochastic mapping using phytools (ARD_phytools(SM)). Results are presented for 15 different generating models and three different tree shapes: symmetrical equal branch length (**A**); asymmetrical ultrametric (**B**); and asymmetrical equal branch length (**C**). Each data point represents the mean of 500 simulated characters. Error bars show the 95% confidence interval of the mean. The corHMM implementation is consistently least erroneous.*

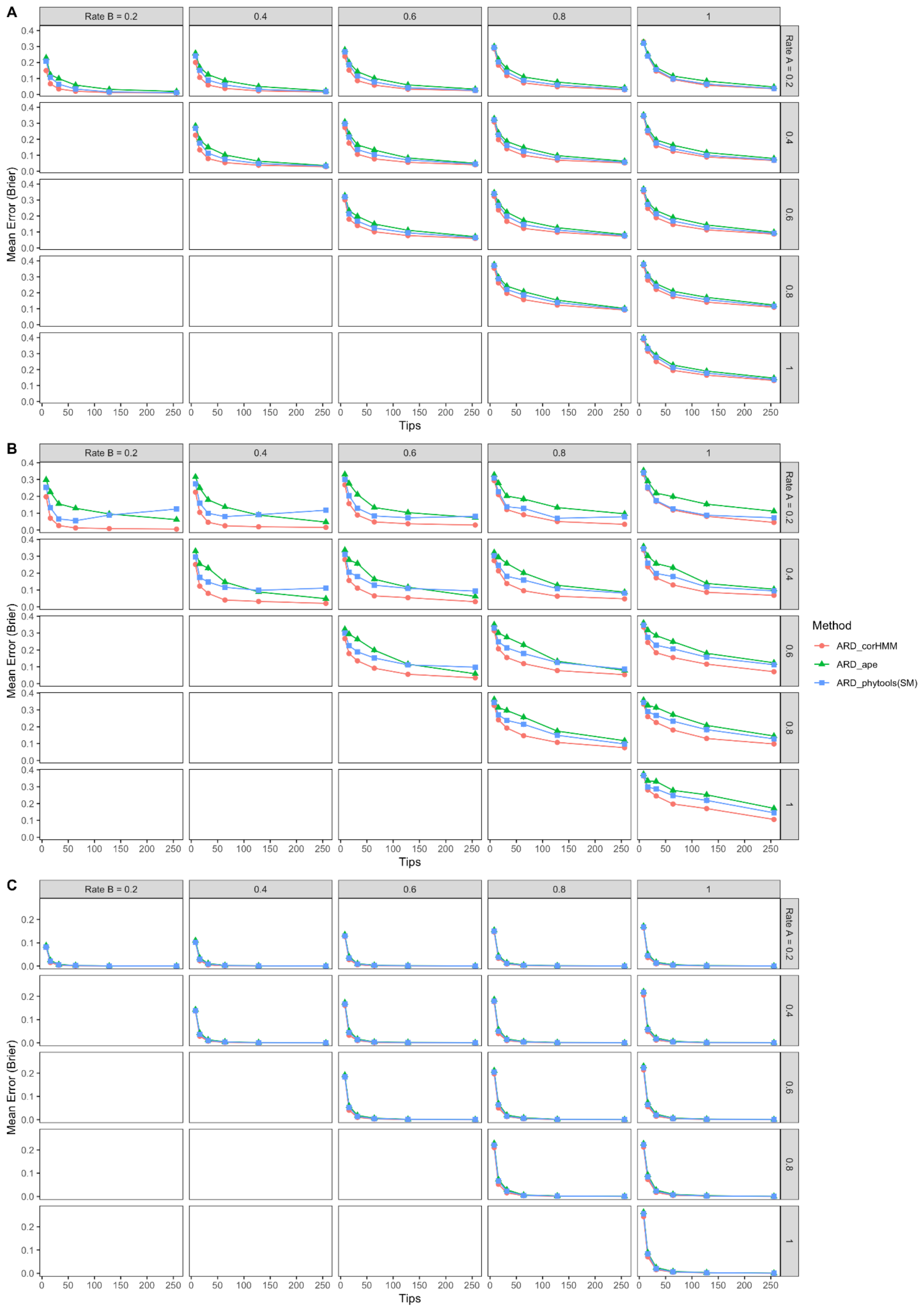


***S. Figure 20.*** *Mean Brier error of the ARD model implemented using different likelihood optimisation methods plotted against the number of tips present in the tree used to simulate character data: Marginal likelihood estimation using corHMM (ARD_corHMM); scaled*

*conditional likelihood using ape (ARD_ape); stochastic mapping using phytools (ARD_phytools(SM)). Results are presented for 15 different generating models and three different tree shapes: symmetrical equal branch length (**A**); asymmetrical ultrametric (**B**); and asymmetrical equal branch length (**C**). Each data point represents the mean of 500 simulated characters. Error bars show the 95% confidence interval of the mean. The corHMM implementation is consistently least erroneous.*

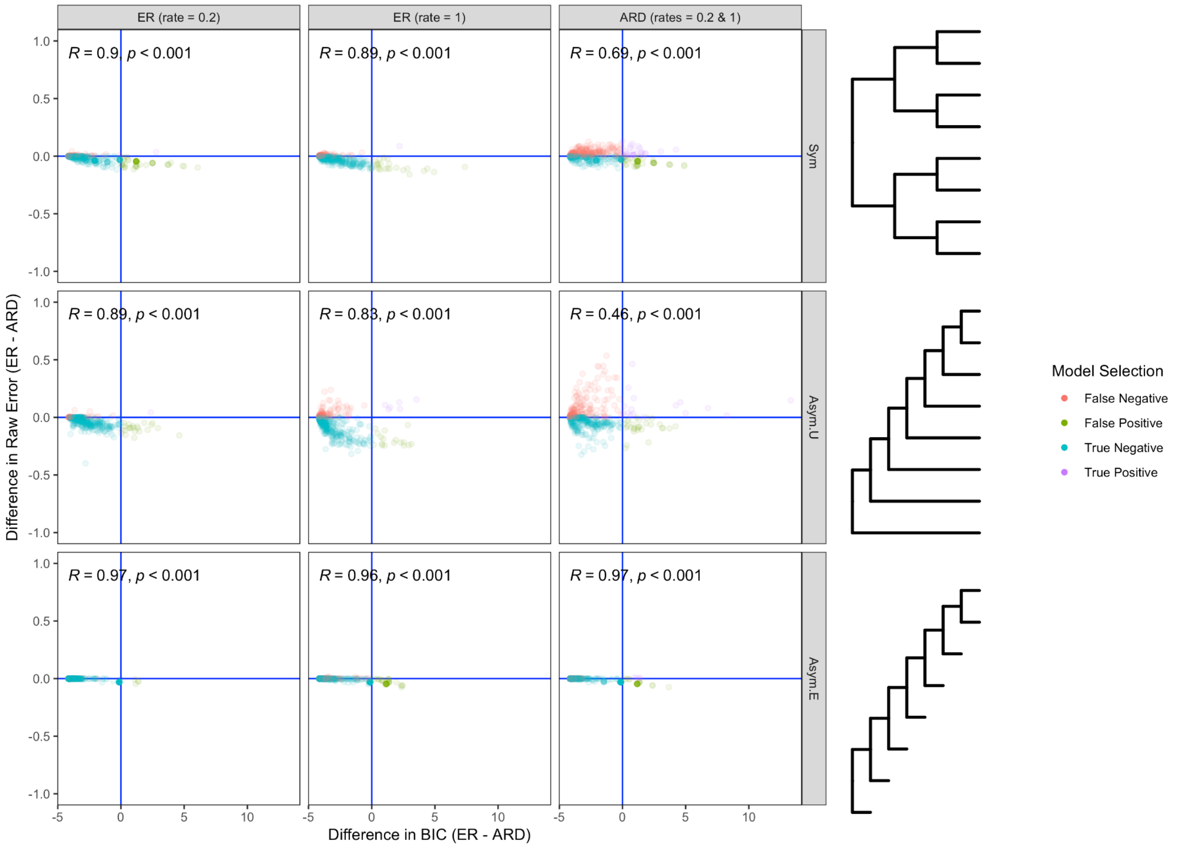


***S. Figure 21.*** *Difference in BIC score between ER and ARD models (ΔBIC) plotted against the difference in raw error between ER and ARD models (Δraw error). Results are presented for data generated under three different models: Equal Rates (rate = 0.2); Equal Rates (rate = 1); All-Rates-Different (rates = 0.2 & 1) using 64-tip trees of three different shapes: symmetrical equal branch lengths (Sym); asymmetrical ultrametric (Asym.U); and asymmetrical equal branch lengths (Asym.E). Top = ER has more error; Bottom = ARD has more error; Left = ER has better BIC score; Right = ARD has better BIC score.*
